# A single cell view of the transcriptome during lateral root initiation in *Arabidopsis thaliana*

**DOI:** 10.1101/2020.10.02.324327

**Authors:** Hardik P. Gala, Amy Lanctot, Ken Jean-Baptiste, Sarah Guiziou, Jonah C. Chu, Joseph E. Zemke, Wesley George, Christine Queitsch, Josh T. Cuperus, Jennifer L. Nemhauser

## Abstract

Root architecture is a major determinant of fitness, and is under constant modification in response to favorable and unfavorable environmental stimuli. Beyond impacts on the primary root, the environment can alter the position, spacing, density and length of secondary or lateral roots. Lateral root development is among the best-studied examples of plant organogenesis, yet there are still many unanswered questions about its earliest steps. Among the challenges faced in capturing these first molecular events is the fact that this process occurs in a small number of cells with unpredictable timing. Single-cell sequencing methods afford the opportunity to isolate the specific transcriptional changes occurring in cells undergoing this fate transition. Using this approach, we successfully captured the transcriptomes of initiating lateral root primordia, and discovered many previously unreported upregulated genes associated with this process. We developed a method to selectively repress target gene transcription in the xylem pole pericycle cells where lateral roots originate, and demonstrated that expression of several of these targets was required for normal root development. We also discovered novel subpopulations of cells in the pericycle and endodermal cell files that respond to lateral root initiation, highlighting the coordination across cell files required for this fate transition.

**One sentence summary:** Single cell RNA sequencing reveals new molecular details about lateral root initiation, including the transcriptional impacts of the primordia on bordering cells.

## Introduction

Plants grow new tissues and organs throughout their lives. To enable this iterative body plan, cells must maintain mechanisms of organogenesis, proliferation, and differentiation. Iterative growth of plants is most easily observed above ground, as plants put out new branches, leaves, and flowers, but equally important is the growth of root systems below the soil. Lateral root development is essential for a plant to remain stably rooted as well as to obtain essential water and nutrients from its surrounding environment. Plants use plasticity in their growth patterns to avoid adverse stimuli and take advantage of favorable ones. In this way, changing root architecture is one of the main mechanisms by which plants can adapt to changing environmental conditions (Khan et al., 2016).

Lateral root development proceeds through three discrete stages, specification, initiation, and emergence. Auxin response exhibits cyclic maxima in the basal meristem with a periodicity of about six hours (De Smet et al., 2007). During specification, cells in the meristematic zone of the primary root are specified as competent to form lateral roots if they transit through the basal meristem when auxin response is high (De Smet et al., 2007). Many other genes oscillate in phase with auxin response in the basal meristem—these genes may be targets of auxin signaling or independent regulators of specification (Moreno-Risueno et al., 2010). These competent cells then exhibit a sustained auxin maximum in the differentiated zone of the root, termed prebranch sites. In *Arabidopsis* and most dicot plants the pericycle cell layer within these prebranch sites is the cell layer that undergoes lateral root initiation (Beeckman et al., 2001). How this initial transient auxin response is molecularly translated to the sustained auxin response of prebranch sites leading to initiation is unknown.

The earliest morphological signal of lateral root initiation is the nuclear migration of two longitudinally adjacent pericycle cells to their shared cell wall. These cells consequently undergo the first anticlinal cell division that initiates lateral root development. *GATA TRANSCRIPTION FACTOR 23 (GATA23)* is necessary for this nuclear migration to occur (De Rybel et al., 2010). *LOB DOMAIN-CONTAINING PROTEIN 16 (LBD16)* and *LOB DOMAIN-CONTAINING PROTEIN 29 (LBD29)* are two other transcription factors shown to play a role in lateral root initiation. These genes are direct targets of *AUXIN RESPONSE FACTOR 7 (ARF7)* and *AUXIN RESPONSE FACTOR 19 (ARF19)* that promote cell division (Okushima et al., 2007). Mutants of these genes exhibit a loss of lateral root initiation, and overexpression of LBD16 rescues the *arf7arf19* mutant phenotype which also lacks lateral roots (Goh et al., 2012). Further cell divisions of specific plane orientations and structural changes of cell files exterior to the pericycle allows for lateral root emergence. The emergence process is accompanied by strong upregulation of cell wall remodelers that are also targets of auxin signaling (Lewis et al., 2013; Ramakrishna et al., 2019) and appears to be the easiest stage of lateral root development to arrest, with many mutants arresting at this stage.

Transcriptomic analyses of lateral root development have been a rich resource for determining key regulators of this developmental process (Vanneste et al., 2005). Careful temporal staging and analyses of different steps during lateral root formation have led to identification of novel regulators (Voß et al., 2015) though the complexity of the pathways regulating this process has also become more apparent. Complicating this analysis is the fact that lateral root development is not cell-autonomous, with many different cell types playing different roles and activating diverse genetic networks during this process. Cell sorting analyses on lateral root development have not been done to parse tissue-specific signals, likely because the regions of the root undergoing this fate transition are prohibitively small for such analyses. Single-cell RNA-sequencing is an alternative approach to obtain transcriptomes on the level of individual cells that requires much less tissue compared to cell sorting. In plants, single-cell RNA-sequencing has been used to characterize several plant tissue types, and single-cell analyses of root transcriptomes have identified both previously characterized and novel cell type markers (Jean-Baptiste et al., 2019; Ryu et al., 2019; Shulse et al., 2019; Shahan et al., 2020). To date, single-cell analyses of root tissue have focused on gene expression in the primary root, transcriptome changes between hair cells and non-hair cells, endodermal differentiation, and regeneration of the primary root meristem after injury.

While initiation of lateral roots is known to be regulated by auxin, only a handful of specific molecular markers of this fate switch have been identified. One reason for this scarcity of markers may be that for any given primary root, lateral root initiation only occurs in a very small proportion of xylem pole pericycle (XPP) cells at near basal meristem, which themselves are a very small proportion of the root cells (approximately five percent) (Schmidt et al., 2014). Rarity of lateral root fate transition is further complicated by the pulsatile nature of the auxin signal, making this a highly transient event (Moreno-Risueno et al., 2010). To counteract these challenges, we microdissected sections of *Arabidopsis* roots undergoing gravity-induced lateral root initiation, and subjected the resulting protoplasts to single-cell sequencing. Using this approach, we successfully captured cells from all major cell types of the root outside the meristem. Through pseudotime analyses found that cells identified as lateral root primordia (LRP) are transcriptionally derived from those identified as xylem pole pericycle cells (XPP), consistent with previous morphological analysis (Malamy and Benfey, 1997). Differential gene analyses identified many previously unreported genes that are upregulated in LRP cells as compared to XPP cells. We validated the expression patterns of a subset of these genes using fluorescent reporters. In addition, we developed a CRISPR/dCas9 tool to specifically target the repression of these candidate genes in XPP cells and found that many of these targets shape root architecture. Finally, we were able to harness the single-cell approach to determine how cells surrounding the developing primordium, specifically endodermal cells overlaying and pericycle cells flanking LRPs, are affected by this fate transition.

## Results

To examine the developmental transition of lateral root initiation, we used gravistimulation to synchronize the formation of lateral root primordia, and then dissected the region of interest at two time points and performed single-cell transcriptome analyses. Mechanical or gravitropic bending of primary roots in *Arabidopsis* causes the accumulation of auxin and the formation of a lateral root at the bend (**Figure 1A**) (De Smet et al., 2007; Ditengou et al., 2008) In our conditions, wild-type plants have formed a primordium at either stage I or II by twenty hours after gravistimulation (Guseman et al., 2015) As our goal was to identify early regulators of lateral root initiation, we analyzed cells twenty hours post-bending, when initiation has just begun, and eight hours post-bending, where there are no morphological signs of lateral root development but transcriptome changes have started (Voß et al., 2015). We included a control treatment group where we did not bend the roots but cut a similar region of the primary root. We microdissected the root bend regions to maximize our yield of the rare cell types of interest.

**Figure 1:**
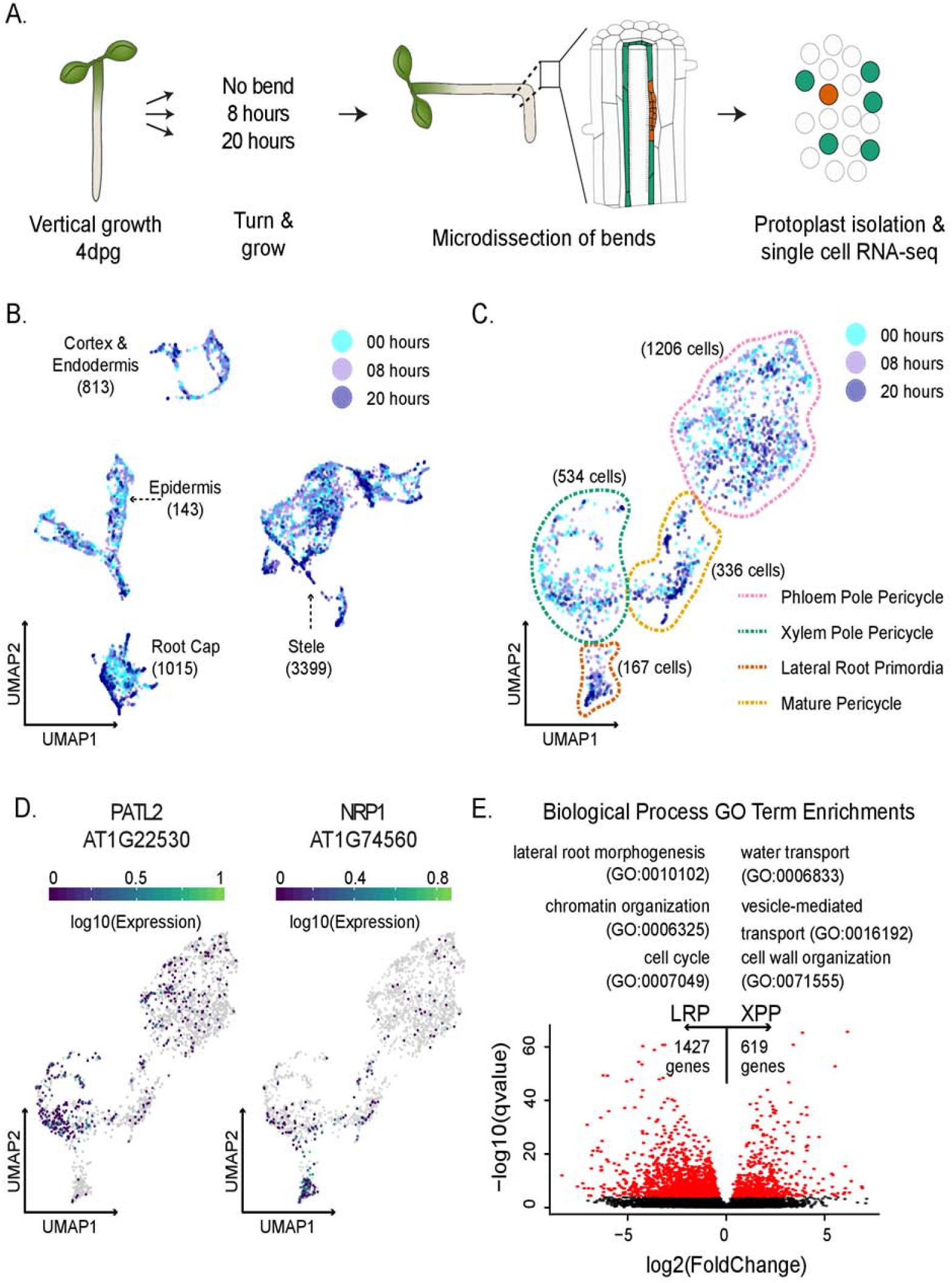
Analysis of lateral root initiation using single-cell RNA-sequencing. A. Experimental design: *Arabidopsis thaliana* seedlings were grown vertically for 4 days post-germination (dpg) then rotated (or marked, in case of control) and grown for an additional eight or twenty hours. Protoplasts were prepared from microdissected root sections for single-cell RNA sequencing. B. UMAP of all 6658 cells colored by experiment. Cell type identities were assigned to each partition based on a set of marker genes. The cell type identities are indicated on the UMAP with the number of cells corresponding to each cell type. C. UMAP of pericycle specific cells, such as lateral root primordia (LRP), mature pericycle (MP), phloem pole pericycle (PPP), and xylem pole pericycle cells (XPP) colored by experiment. D. Expression pericycle-specific UMAPs of two DEGs between LRP and XPP. E. Scatter plot of the log2 fold change of average gene expression between XPP and LRP and the Mann Whitney Wilcoxon qvalue. DEGs are colored in red. A selection of GO Terms associated with DEGs are shown above the plot.

Our experiment yielded 6658 cells with a mean number of 15987 reads per cell and a median of 1383 genes expressed. Of these 6658 cells, 1730 (~26%) cells were from the twenty-hour timepoint, 2443 (~37%) cells were from the eight-hour timepoint, and 2485 (~37%) cells were from the no bend control (**Supplemental Table 1**). Further analysis was performed using Monocle 3 (Trapnell et al., 2014; Qiu et al., 2017; Cao et al., 2019), which uses Principal Component Analysis (PCA) and Uniform Manifold Approximation and Projection (UMAP) to reduce the dimensionality of the dataset and to visualize the relationships among cellular transcriptomes in a two-dimensional space. Our analysis revealed five clusters. Using previously defined cell type markers (Brady et al., 2007; Cartwright et al., 2009), each cluster was assigned a label. Of the 6658 cells, 813 (~12%) cells were classified as cortex & endodermis, 1015 (~15%) cells were classified as columella/root cap, 1431 (~22%) cells were classified as epidermis, and 3399 (~51%) cells were classified as stele (**Figure 1B, Supplemental Figure 1A**). These results show that the microdissection was successful in capturing a representative sample of stele cells, as the proportion of these cells in our data was similar to that determined by imaging analyses (Shulse et al., 2019).

### Xylem pole pericycle cells are precursors of mature pericycle and lateral root primordia cells

To better understand the relationships between vasculature cell types, we re-clustered all cells identified as belonging to the stele. Of the 3399 stele cells, 216 cells were labeled phloem, 242 cells were labeled xylem, 1206 cells were labeled phloem pole pericycle (PPP), 534 cells were labeled XPP, 336 cells were labeled mature pericycle, 167 cells were LRP, and 698 cells were too ambiguous in their gene expression to assign a label (**Supplemental Figure 1B, Supplemental Figure 1C**, **Supplemental Table 1**). A pericycle-specific UMAP was also generated that included XPP, LRP, PPP and mature pericycle cells (**Figure 1C**). A cell developmental trajectory connected the pericycle sub-clusters and recapitulated the known developmental relationship between these cell types. Trajectories initiating either from the XPP or the PPP converged to mature pericycle but only the XPP (and not the PPP) population branched out toward the LRP, confirming that our results were faithfully recapitulating the known exclusive relationship of XPP to LRP cells (Malamy and Benfey, 1997). The XPP and the PPP cells are expected to differentiate at a similar time, whereas LRP and mature pericycle cells should form later. Our dataset confirmed this developmental progression, as approximately twenty percent of the XPP and PPP cell clusters were from the twenty-hour timepoint whereas approximately fifty percent of the LRP and mature pericycle clusters were from this later timepoint. (**Supplemental Figure 1D**).

We next attempted to identify genes expressed as XPP cells transitioned to LRP cells. To do this, we carried out a pseudotime analysis originating from the XPP population and connecting to either the LRP or the mature pericycle (**Supplemental Figure 2A, Supplemental Figure 2B**). Using a false discovery rate (FDR) cutoff of 0.1, 1892 genes were identified as changing in at least one of the trajectories. Of these genes, 878 genes were specific to the XPP to mature pericycle trajectory, 504 genes were specific to the XPP to LRP trajectory, and 510 genes were observed in both trajectories (**Supplemental Figure 2C, Supplemental Figure 2D**). As XPP cells develop, expression of genes found in both trajectories decreased and most were not expressed at high levels in the LRP cells or the mature pericycle cells. Genes uniquely upregulated across the XPP to the LRP trajectory were overrepresented with transcription factors including the well-known markers of initiation *LBD16* and *LBD29* (Okushima et al., 2007) as well as *WUSCHEL-RELATED HOMEOBOX 5* (*WOX5*), *CYTOKININ RESPONSE FACTOR 2* (*CRF2*) and *PUCHI*, all of which have previously been shown to play a role or be expressed in lateral root development (Hu and Xu, 2016; Jeon et al., 2016; Goh et al., 2019) **(Supplemental Figure 2E).** Other genes unique to this trajectory were regulatory kinases like *MUSTACHES* (*MUS*), *MAP KINASE KINASE 6* (*MKK6*), and *RGF INSENSITIVE 5* (*RGI5*), which have also been shown to regulate lateral root development (Zeng et al., 2011; Ou et al., 2016; Xun et al., 2020) **(Supplemental Figure 2E)**.

We next performed differentially expressed genes (DEG) analysis on the transcriptomes associated with these different populations. Due to the small number of LRP cells (167 cells i.e. ~2.5% of all cells), three different statistical approaches were used to perform DEG analysis: a generalized linear model (GLM), the Mann Whitney Wilcoxon (MMW) test, and a recently published packaged called Vision (DeTomaso et al., 2019). We used previously identified LRP-enriched genes as a guide to inform our use of the results from each method. *LBD16* and *LBD29* were called in all three methods, whereas *ARF19* and *GATA23* were only called in Vision (but narrowly missed with p-values of 0.001 and 0.0004 respectively in MWW method). ARF7 was not significantly different in any approach. To compile the most comprehensive list, we generated a list of DEGs that were significantly different between XPP and LRP populations in at least two approaches. We called 1427 DEGs specific to LRP and 619 DEGs specific to XPP cells. Several of these genes have been previously characterized as specific in their expression patterns, and expression maps of these genes reflect this quality (**Figure 1D, Supplemental Figure S3, Supplemental Data 1**) (Zhu et al., 2006; Tejos et al., 2018).

As expected, a Gene Ontology (GO) term enrichment analysis of the DEGs with higher expression in LRP cells showed a strong enrichment for terms associated with lateral root formation, lateral root morphogenesis, lateral root development, and auxin response. Terms associated with the regulation of translation initiation and RNA processing (**Supplemental Data 1**) were also enriched, indicating that the transition from the XPP to the LRP requires a burst of *de novo* protein production. Increased protein production is associated with a stem cell state (Himanen et al., 2004), which correlates with the transition from XPP to early LRP cells which are competent to form all the types of cells of a developing root. Other GO terms associated with the DEGs more highly expressed in LRP were lateral root morphogenesis, cell cycle, and chromatin organization (**Figure 1E**).

### Single-cell analysis recapitulates and extends findings from previous transcriptome studies

Lateral root development has been extensively characterized with transcriptomic analysis, generated using a variety of lateral root induction models and experimental designs. We compared our LRP versus XPP DEGs with published datasets to assess differences in single-cell versus whole tissue (or population) methods. We found that roughly half of the genes (including genes regulated by auxin and genes belonging to cell cycle processes) identified by a microarray analysis of induced lateral root development (Vanneste et al., 2005) were included in our set of genes upregulated in LRP cells; only five of these genes were in our set of XPP-upregulated genes (**Supplemental Figure S4A**). This result suggests our approach faithfully captured genes involved in lateral root initiation and distinguished LRP and XPP cells. A detailed time course analysis of lateral root development using a similar bend assay as what we applied (Voß et al., 2015) allowed us to compare to bulk RNA sequencing data taken at similar timepoints as used in our study. Two thirds of the genes previously found upregulated at nine hours post-bending were contained within our set of genes upregulated in LRP, none of these genes were XPP-upregulated (**Supplemental Figure S4B**). Roughly half the genes found upregulated at twenty-one hours were within our LRP-upregulated gene list; only nine were in our XPP-upregulated gene list (**Supplemental Figure S4B**). At both time points, approximately one thousand genes were uniquely found in LRP DEGs from our study. The genes that were identified in the previous study but not in ours may be attributed to technical differences or likely are expressed in cell layers outside the pericycle.

We also compared our data to transcriptome assays that did not directly examine lateral root initiation through root bending, but queried related processes in the root. We compared our data to a time course analysis of primary root transcriptomes after auxin treatment (Lewis et al., 2013), as auxin treatment strongly promotes lateral root initiation. We found that thirty-seven LRP-upregulated genes were strongly induced in this dataset in response to auxin, whereas only two XPP-upregulated genes were auxin-induced (**Supplemental Figure S4C**). In contrast, twenty-six XPP-upregulated genes were repressed by auxin treatment, whereas only five LRP-upregulated genes were in this repressed dataset (**Supplemental Figure S4C**). Another recent analysis identified genes specifically induced by ARF19-mediated auxin response (Powers et al., 2019). We found that 243 of our LRP-upregulated genes overlapped with the set of ARF19-specific auxin-induced genes, while only 19 XPP-upregulated genes did so (**Supplemental Figure S4D)**. We conclude that our data reflect the auxin-inducibility of lateral root initiation, and specifically that this auxin inducibility was at least in part mediated by ARF19. ARF19 is unique among the ARFs in being both auxin-responsive in its own expression pattern (Wilmoth et al., 2005) and a very strong activator of transcription itself (Lanctot et al., 2020)

Finally, we examined how our dataset compared to genes expressed in the basal meristem during lateral root specification. During specification, cells become competent to form lateral roots if they transit through the basal meristem during an auxin response maximum. Many genes exhibit similar oscillatory behavior to auxin response in the basal meristem (Moreno-Risueno et al., 2010). We found that fifty-eight genes that oscillated in phase with auxin response in the basal meristem were in our set of LRP-upregulated genes, whereas only one XPP-upregulated gene oscillated in phase with auxin (**Supplemental Figure S4E**). However, 213 XPP-upregulated genes oscillated antiphase to auxin in the basal meristem, while only twenty LRP-upregulated genes show antiphase oscillation. How specification and initiation are connected temporally and spatially and how competent cells “remember” their future cell fate is still unknown. Our results suggested that oscillatory behavior of some genes may predict their importance during initiation later in development, and in particular that genes with antiphase oscillation patterns may actively repress lateral root fate. We also compared our dataset to a study that determined genes whose expression was impacted by repressing auxin response specifically in early-stage LRPs (Ramakrishna et al., 2019), and found our LRP-upregulated genes overlapped with nearly two hundred of these genes (**Supplemental Figure S4F**), again emphasizing the importance of auxin response for establishing lateral root fate.

### Genes upregulated in LRP cells are indicative of cells undergoing fate transitions

We selected timepoints that should best capture the transition of undifferentiated XPP cells into LRPs. As one of the earliest morphological steps in this process is an asymmetric cell division, followed by many subsequent cell divisions, it is not surprising that a number of LRP-enriched DEGs are involved in cell cycle control. It is already known, for example, that expression of cyclin *CYCB1;1* marks lateral root primordia (Beeckman et al., 2001) Interestingly, one of the distinguishing features of XPP cells when compared to PPP cells is that some XPP cells are arrested in G2, whereas all PPP cells are arrested in G1 (Beeckman et al., 2001). This G2 arrest may prime these XPP cells to undergo rapid reintroduction into the cell cycle. We also found enrichment of chromatin remodeling factors in the set of LRP-upregulated genes. Considering lateral root initiation is the first step of organogenesis, it makes sense that broad transcriptional changes, mediated by changes in the chromatin landscape, may be required, but this has not previously been reported (**Supplemental Data 1**). Finally, we found that genes that promote cell division, differentiation, and “stemness”, mostly transcription factors, were also enriched in the set of genes upregulated in LRP cells. Most of these genes have been characterized as regulating development in other meristems, such as the primary root meristem or the shoot apical meristem, but had not been shown to play a role in lateral root development. We chose several candidate genes from these three groups, chromatin regulators (four genes), cell cycle regulators (seven genes) and stemness regulators (six genes) to carry out two types of validation experiments: (1) characterizing their spatial expression patterns with transcriptional reporters in wild-type Col-0 and *arf7arf19* mutant seedlings, and (2) phenotypic evaluation of lateral root development in transgenic line with cell-type specific repression of candidate genes.

### A novel cell-type specific dCas9-driven repressor system can reveal drivers of lateral root fate

To explore the functional role of candidate genes in lateral root development, we devised a method to repress candidate genes only in the XPP cell lineage. We leveraged the enhancer trap line J0121 which is specifically expressed in XPP cells via a UAS-GAL4 driver system (Laplaze et al., 2005). We first introgressed J0121 into the Col-0 background (referred to hereafter as J0121^Col^), and then introduced a UAS-dCas9-TPLN300 repressor (dCas9R) construct with three gene-specific sgRNAs directed to the promoter regions of candidate genes (J0121^Col^≫dCas9R, **Supplemental Figure S5, Supplemental Data 2**). This cell-type specific repression system has several advantages over traditional knockdown and knockout studies. For instance, multiple guides can be used to simultaneously repress several members of the same gene family that may have redundant functions. Additionally, many of the candidate genes we identified as enriched in LRP cells also play roles in embryonic and primary root development, greatly complicating assessment of any role in lateral roots. To test the efficacy of our assay, we tested the effects of repressing expression of both *ARF7* and *ARF19*. We found that these perturbation lines have significantly reduced lateral root density compared to the empty vector control as expected although they did not fully recapitulate the full suppression of lateral roots seen in *arf7arf19* null mutants (**Supplemental Figure S6, Figure 2C**). This observation is consistent with the small reduction of lateral root number seen in a GATA23-driven CRISPR/Cas9 deletion of *ARF7* and *ARF19* (Decaestecker et al., 2019). There are at least two likely explanations for the milder phenotype of J0121^Col^≫dCas9R compared with the *arf7arf19* null mutant. First, repression in J0121^Col^ expression is limited to XPPs and lateral root stages I through III, so there is likely residual ARF protein that persists from expression in pre-XPP fate cells (Dubrovsky et al., 2006). Second, our synthetic repressor may not block all transcriptional activity, leading to hypermorphic rather than amorphic phenotypes. Even given these limitations, the system proved sufficiently sensitive to enable detection of cell type-specific impacts on lateral root development.

### Chromatin remodeling factors influence lateral root development

Three histone deacetylases (HDACs) that are all in the same plant-specific gene family, *HISTONE DEACETYLASE 3, HISTONE DEACETYLASE 2, and HISTONE DEACETYLASE 13 (HDA3, HD2B, and HDT4*) (Li et al., 2017; Luo et al., 2017), were all enriched in LRP cells in our DEG analysis **(Figure 2A)**, as was the E3 ubiquitin ligase *ORTHRUS 1 (ORTH1*) which decreases DNA methylation (Kim et al., 2014). Transcriptional reporters of these genes express strongly in early stage primordia **(Figure 2B)**. Expression of *HDT4* and *HDA3* were specific to LRPs in the differentiated zone of the primary root, though both were also strongly expressed in the meristematic zone of the primary root **(Supplemental Figure S7)**. Their expression in the meristem was strongly decreased in *arf7arf19* mutants, suggesting they may be regulated by auxin **(Supplemental Figure S7)**. *HD2B* was also strongly expressed in LRP cells, as well as in the primary root meristem and other pericycle cells (**Supplemental Figure S7**). *ORTH1* was broadly expressed in the vasculature of the differentiated zone of the primary root, not only in LRP cells **(Supplemental Figure S7)**, which is reflected by its enrichment in mature pericycle cells in our single-cell library **(Figure 2A)**. Its expression was not impacted in *arf7arf19* mutant lines **(Supplemental Figure S7)**.

**Figure 2.**
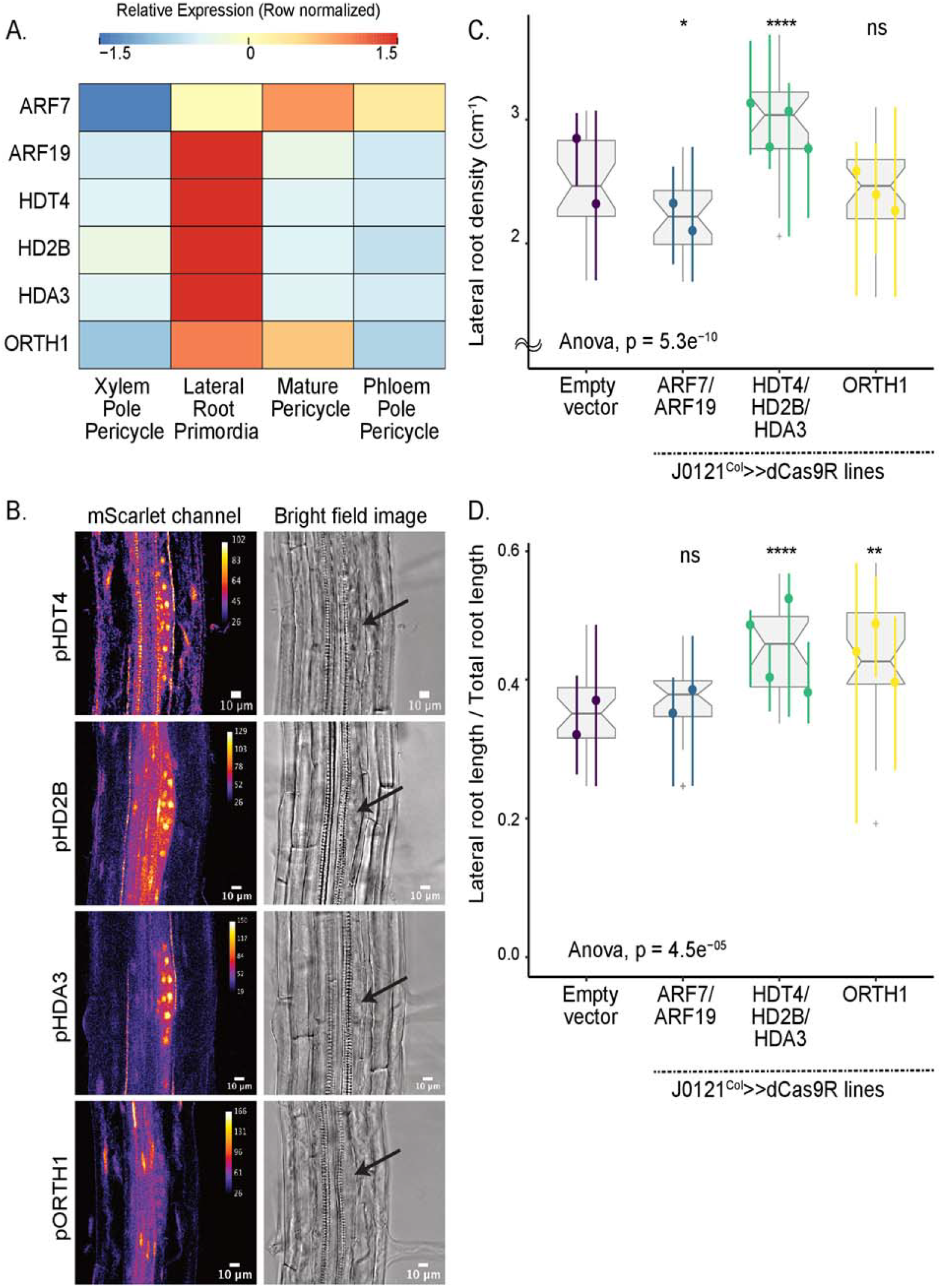
Validation experiments on chromatin modifier candidate genes. A. Heatmap (row-scaled) visualizing expression of candidate genes in the pericycle cell clusters from the single-cell library. Scale bar represents the z-score of the normalized expression values. B. Confocal microscopy images of candidate genes’ transcriptional reporters in early stage lateral root primordia (left) and bright field image of corresponding primordia (right). C. Lateral root density of J0121^Col^≫dCas9R transgenic lines of candidate genes. D. Proportion of total root length contributed by lateral roots of J0121^Col^≫dCas9R transgenic lines of candidate genes. For C and D, significance was determined by pairwise comparison with empty vector control and ANOVA. For each candidate gene, multiple independent transgenic lines have been analyzed; each colored line in the graph represents an individual line.

Using J0121^Col^≫dCas9R, we targeted all three *HDAC* genes for repression using distinct guide RNAs for each gene. Simultaneous repression of all three genes caused a strong phenotype, where both the density of lateral roots **(Figure 2C)** and the proportion of total lateral root length contributed by lateral roots were significantly increased **(Figure 2D)**. The phenotype suggested that the HDACs may repress lateral root initiation and later stages of development. *ORTH1* repression in XPP cells did not significantly impact lateral root density **(Figure 2C),** but the proportion of total root length contributed by lateral roots significantly increased **(Figure 2D)**. Thus, *ORTH1* may repress lateral root growth only post-initiation. All of these chromatin regulators were expressed strongly in LRP cells, and repressing their function stimulates lateral root growth, suggesting they may act to coordinate cells and promote orderly development.

### Cell cycle regulators are active during lateral root development

We characterized the role of five DEGs enriched in our LRP population that play a role in cell cycle regulation: *RECEPTOR FOR ACTIVATED C KINASE 1A, B* and *1C (RACK1A, RACK1B* and *RACK1C*), *NAP1-RELATED PROTEIN 1* and *2 (NRP1 and NRP2)*, and *CYCLIN-DEPENDENT PROTEIN KINASE INHIBITORS 6* and *11 (SMR6* and *SMR11)* **(Figure 3A)**. *RACK1B* and *RACK1C* interact with protein kinase C (Guo and Chen, 2008). *SMR6* and *SMR11* are cyclin-dependent kinase inhibitors (Yi et al., 2014), and *NRP1* is a histone chaperone required for the G2 to M transition (Zhu et al., 2006). We generated transcriptional fluorescent reporters of *RACK1B* and *RACK1C* and found their expression was indeed specific to early stage lateral root primordia as seen in the single-cell data **(Figure 3B)**. Expression of both *RACK1B* and *RACK1C* was lost in the differentiated zone of the primary root in *arf7arf19* mutants, which do not form lateral roots, indicating the specificity of their expression to LRP in this zone **(Supplementary Figure S8)**. Expression of these genes in the primary root meristem also was highly decreased in *arf7arf19* mutants **(Supplementary Figure S8)**, suggesting ARF7 and ARF19 may be the primary ARFs regulating their expression throughout the root.

**Figure 3.**
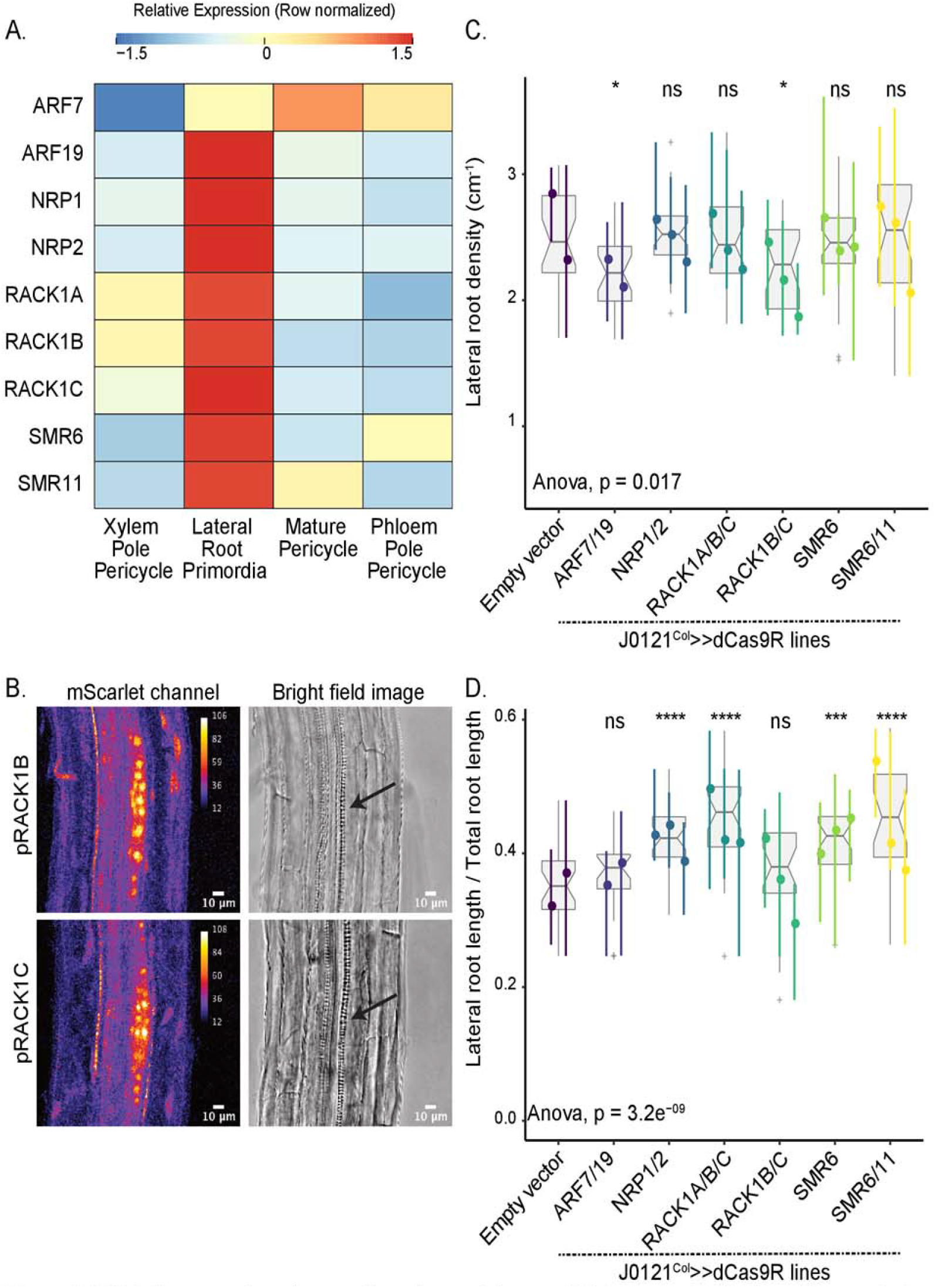
Validation experiments on cell cycle regulator candidate genes. A. Heatmap (row-scaled) visualizing expression of candidate genes in the pericycle cell clusters from the single-cell library. Scale bar represents the z-score of the normalized expression values. B. Confocal microscopy images of candidate genes’ transcriptional reporters in early stage lateral root primordia (left) and bright field image of corresponding primordia (right). C. Lateral root density of J0121^Col^≫dCas9R transgenic lines of candidate genes. D. Proportion of total root length contributed by lateral roots of J0121^Col^≫dCas9R transgenic lines of candidate genes. For C and D, significance was determined by pairwise comparison with empty vector control and ANOVA. For each candidate gene, multiple independent transgenic lines have been analyzed; each colored line in the graph represents an individual line.

We next used J0121^Col^≫dCas9R to test for functional relevance of these cell cycle regulators. *NRP1* is highly related to *NRP2*, so we generated a repression line with guides targeting both genes. These lines did not show differences in lateral root density **(Figure 3C),** but did show a significantly increased proportion of summed total root length that was contributed by lateral roots reflecting longer lateral roots than in the control lines **(Figure 3D)**. *RACK1B* and *RACK1C* are highly related and show redundancy with *RACK1A*, so we generated two separate sets of repression lines, one with guides targeting *RACK1B* and *RACK1C* and the other with guides targeting all three *RACK1* genes. Repression of *RACK1B* and *RACK1C* caused a significant decrease in lateral root density **(Figure 3C)**. Interestingly, while repression of all three *RACK1* genes did not significantly impact lateral root density, this line did show a significantly increased proportion of total root length contributed by lateral roots **(Figure 3D)**, similar to the *NRP1/NRP2* repressed line. Singular repression of *SMR6* and concurrent repression of *SMR6* and *SMR11* did not show differences in lateral root density from the control **(Figure 3C),** but again showed significantly increased proportion of total root length contributed by lateral roots **(Figure 3D)**.

### Genes that encode pluripotency and stemness are upregulated in LRP cells

We chose five DEGs known to play a role in developmental transitions for further validation studies, specifically *TARGET OF MONOPTEROS 6 (TMO6)*, a Dof-type transcription factor originally isolated as a target of ARF5 in embryos (Schlereth et al., 2010), *BREVIS RADIX-LIKE 1 (BRXL1)*, a BRX-like regulator of primary root development (Briggs et al., 2006), *LATERAL ROOT PRIMORDIUM 1 (LRP1)*, which a marker and plays a role in lateral root development (Smith and Fedoroff, 1995; Singh et al., 2020), *OVATE FAMILY PROTEIN 8 (OFP8)*, a transcriptional repressor of KNOX family transcription factors (Wang et al., 2011), and *PLETHORA 3 (PLT3)*, a *PLETHORA* family gene that interprets auxin gradients in the primary root (Santuari et al., 2016). The presence of genes such as *LRP1*, *BRXL1*, and *PLT3*, known to regulate early stages of root development, confirmed that our dataset was isolating genes expected to be active early during lateral root initiation. BRXL proteins have recently been shown to play a role in promoting nuclear migration and asymmetric cell division in the development of stomata (Rowe et al., 2019; Muroyama et al., 2020), a process that is also essential during the very first stages of lateral root initiation. *OFP8* and *TMO6* were somewhat unexpected discoveries, given that ovate family proteins have primarily been characterized in fruit development (Wang et al., 2016) and *TMO* genes have primarily been characterized in embryonic development (Schlereth et al., 2010).

Expression of all of these genes showed enrichment in LRP cells, though *TMO6* expression was most enriched in PPP cells (**Figure 4A**), consistent with its known role in phloem cell division and differentiation (Miyashima et al., 2019). Reporters of *PLT3* (Galinha et al., 2007), *LRP1* (Smith and Fedoroff, 1995) and *TMO6* (Schlereth et al., 2010) have previously been published, so we only generated transcriptional reporters of *OFP8* and *BRXL1*. Both showed strong and specific expression in lateral root primordia (**Figure 4B**). Expression of the *TMO6* reporter had not been previously analyzed in XPP or LRPs. We found that indeed it was strongly expressed in developing primordia, as predicted from the single-cell analysis. Nearly all of this expression was lost in *arf7arf19* mutants (**Supplemental Figure 9**), suggesting that, at least in the root, *TMO6* is primarily a target of these ARFs rather than ARF5. *TMO6*, *BRXL1* and *OFP8* were not expressed in the primary root meristem, (**Supplemental Figure 9**) making them potentially useful for targeting engineering efforts specifically to lateral roots.

**Figure 4.**
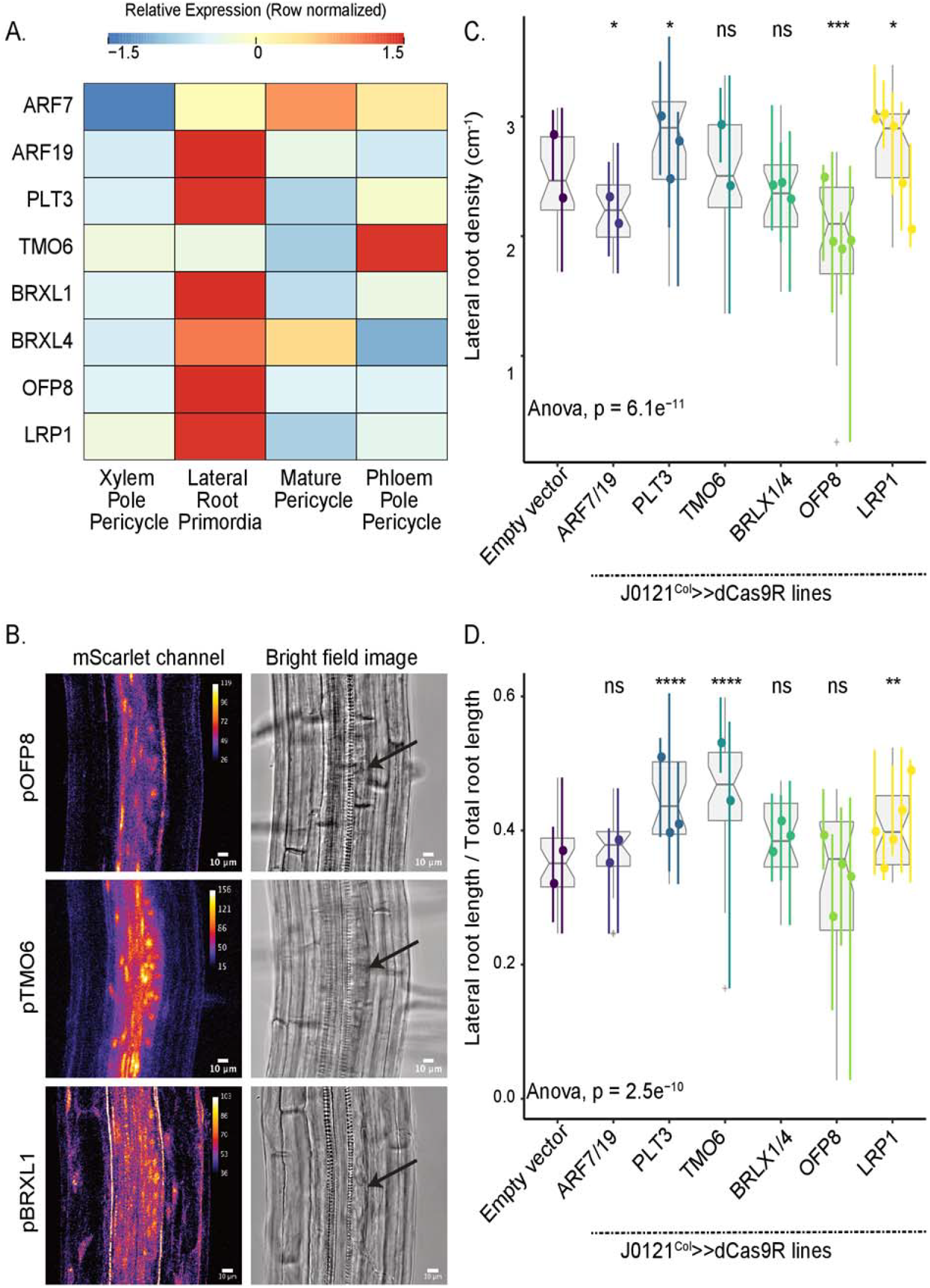
Validation experiments on stemness candidate genes. A. Heatmap (row-scaled) visualizing expression of candidate genes in the pericycle cell clusters from the single-cell library. Scale bar represents the z-score of the normalized expression values. B. Confocal microscopy images of candidate genes’ transcriptional reporters in early stage lateral root primordia (left) and bright field image of corresponding primordia (right). C. Lateral root density of J0121^Col^≫dCas9R transgenic lines of candidate genes. D. Proportion of total root length contributed by lateral roots of J0121^Col^≫dCas9R transgenic lines of candidate genes. For C and D, significance was determined by pairwise comparison with empty vector control and ANOVA. For each candidate gene, multiple independent transgenic lines have been analyzed; each colored line in the graph represents an individual line.

Repression of these genes in XPP cells using J0121^Col^≫dCas9R impacted root architecture but in different ways. Repression of *PLT3* caused a significant increase in both lateral root density (**Figure 4C**) and the proportion of total root length contributed by lateral roots (**Figure 4D**), suggesting *PLT3* may repress both lateral root initiation and emergence post-initiation. Repression of *TMO6* did not impact lateral root density (**Figure 4C**), but did cause a significant increase in the proportion of total root length contributed by lateral roots (**Figure 4D**), suggesting it may act on lateral root development post-initiation.

Repression of *LRP1* in XPP cells significantly increased both lateral root density (**Figure 4C**) and the proportion of total root length that are lateral roots (**Figure 4D**), a phenotype that matches previously reported overexpression lines of *LRP1*, which showed decreased lateral root density (Singh et al., 2020). Concurrent repression of *BRXL1* and its close homolog *BRXL4* did not significantly impact either lateral root density (**Figure 4C**) or the proportion of total root length that are lateral roots (**Figure 4D**), despite its strong expression in LRP cells. We did observe irregular spacing of lateral roots and shorter primary roots in *BRXL/BRXL4* repression lines, suggesting they may play a role in lateral root emergence (**Supplemental Figure 9**). Repression of *OFP8* exhibited unique behavior in our perturbation lines, as these lines showed significantly decreased lateral root density (**Figure 4C**). *OFP8* has not previously been characterized to play any role in root development, and this strong effect on lateral root initiation and its strongly specific expression in LRP is notable. *OFP8* repression did not impact the proportion of total root length contributed by lateral roots (**Figure 4D**) but these lines had shorter primary roots (**Supplemental Figure 6**).

### Non-LRP cells populations undergo transcriptional changes and fate transitions in response to lateral root initiation

Formation of a new lateral root is a self-organizing process during which a very limited number of competent XPP cells undergo repeated cell divisions to initiate lateral root organogenesis (Torres-Martínez et al., 2020). Continued development of the new root requires biophysical restructuring of the surrounding cell files. Signatures of lateral root development are seen outside the pericycle at pre-emergence stages of development, including during early initiation (Vermeer et al., 2014). Feedback on auxin signaling and changes in auxin transport patterns in the endodermis (Marhavý et al., 2013) and the vasculature (De Smet et al., 2007; Porco et al., 2016) are also essential for the first steps of lateral root initiation. Because our single-cell RNA sequencing dataset allowed us to examine the transcriptional state of these different cell layers independently, we examined which of the non-pericycle cell files contribute to transcriptional changes in response to this fate switch. For this we leveraged DEGs (945 genes) identified from a previous bulk RNA transcriptome study of bend-induced lateral root initiation (Voß et al., 2015) corresponding to the twenty hour post-bend timepoint in our study, and mapped the expression of these genes to our cell type resolved dataset **(Figure 5A)**. As expected, most of these genes showed high expression in the LRP population and very low expression in the XPP population. In addition, we found strong enrichment of certain groups of genes in non-LRP populations, especially in those categorized as mature pericycle, endodermis cells and root cap cells.

**Figure 5.**
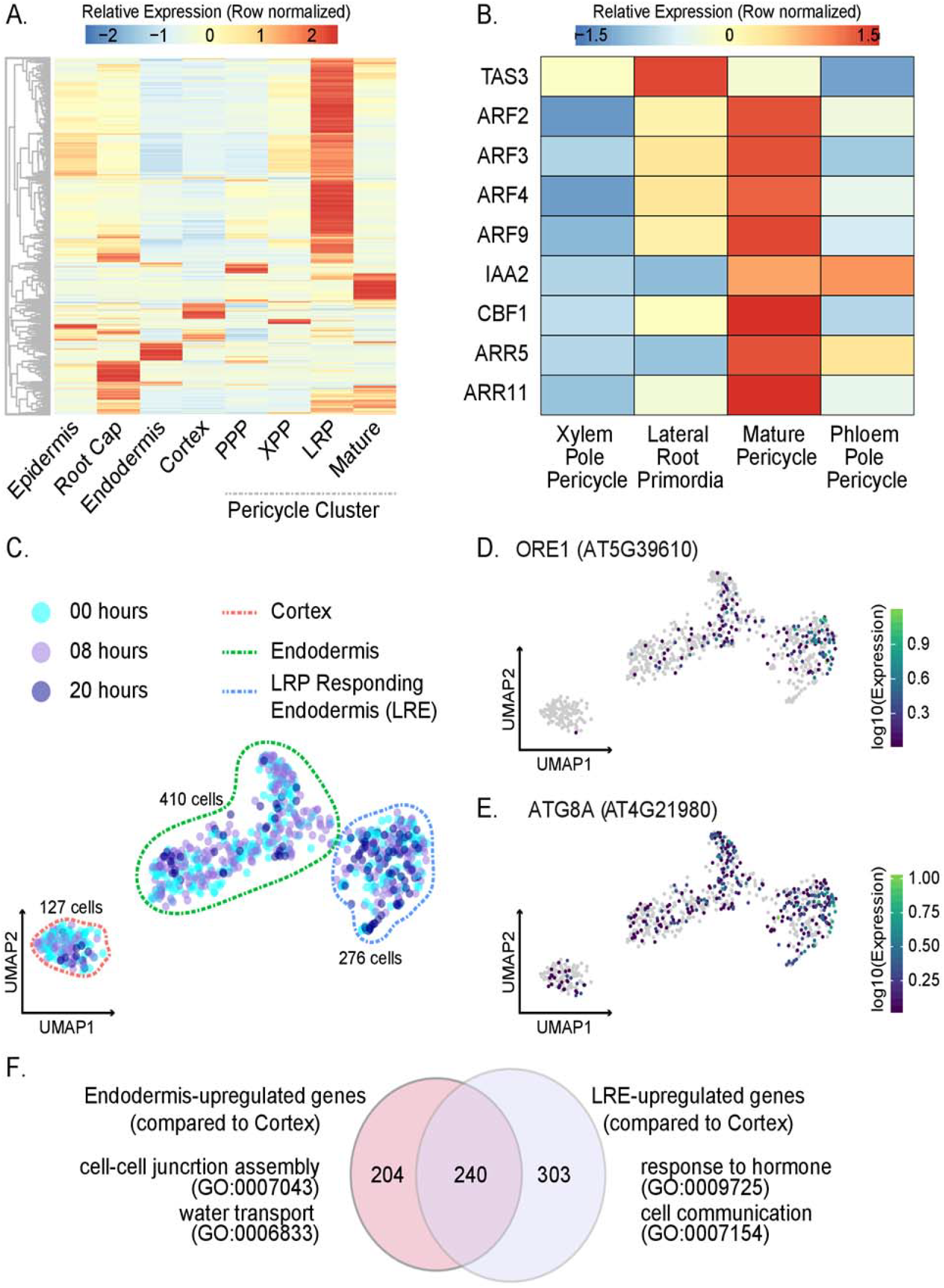
Analysis of non-LRP cells in the single-cell library. A. Heatmap (row-scaled) of expression patterns of previously-identified LR-specifying genes in different cell clusters. B. Expression heatmap (row-scaled) of specific auxin-inducing and auxin-repressing genes in different stele cell clusters. C. UMAP of Cortex, Endodermis, and Lateral Root Responding Endodermis (LRE) cells. D. Expression UMAP of ORE1, a NAC transcription factor that promotes autophagy of endodermal cells overlying LRPs. E. Expression UMAP of AT8GA. F. Gene ontology of genes enriched in endodermis and LRE cell populations. The Venn diagram represents DEGs upregulated in endodermis in comparison to the cortex and upregulated in LRE in comparison to the cortex. A selection of GO terms for each of these sets of DEGs are shown.

We focused first on genes showing strong expression in the mature pericycle, as these cells could be directly in contact with the initiating primordium. To assess if the genetic circuits regulating the XPP to mature pericycle transition were linked to lateral root development at the bend, we examined the genes expressed in our pseudotime analysis of the transition of XPP cells to mature pericycle cells as compared to the genes expressed in the trajectory the transition of XPP cells to LRP cells (**Supplemental Figure 2A-D**). Genetic signatures in this mature pericycle trajectory suggested that these cells, acting as the outgroup for this fate transition, nonetheless were responding to lateral root development, though in a very distinct way from cells within the XPP to LRP trajectory. One gene that was expressed in XPP cells and LRP cells close to the branch point is *TRANS-ACTING siRNA3* (*TAS3)*. *TAS3* promotes lateral root development by inhibiting the class B auxin response factors *ARF2*, *ARF3* and *ARF4* (Marin et al., 2010) Consistent with this pattern, *ARF2* transcripts were found only in the mature pericycle trajectory. While *TAS3* expression is fairly consistent across XPP, LRP, and mature pericycle cells, *ARF2*, *ARF3*, and *ARF4* show greater expression in the mature pericycle cell population as compared to the XPP population, and also show high expression in PPP cells (**Figure 5B**). As LRPs never initiate from PPP, strong expression of these repressor ARFs in mature pericycle cells and PPP cells may indicate that these genes are repressing auxin response, and thus lateral root development, in these cells.

A growing body of research has shown that LRPs send signals to actively repress initiation of surrounding pericycle cells, allowing for the proper spacing of lateral roots along the axis of the primary root (Murphy et al., 2016; Toyokura et al., 2019; Trinh et al., 2019). The strong expression of genes that repress auxin response in mature pericycle cells suggests one mechanism for this inhibition. Further evidence in support of this model is the expression of another repressor ARF, *ARF9*, and the auxin repressor *IAA2* only in the mature pericycle trajectory. Uniquely expressed in this trajectory are also two cytokinin response factors, *ARABIDOPSIS THALIANA RESPONSE REGULATORS 5* and *11 (ARR5* and *ARR11)*, which are known to inhibit auxin response, repress lateral root development and coordinate uniform spacing of lateral roots (Mason et al., 2005; To et al., 2004). A target of cytokinin signaling, *CFB1*, which is specifically expressed in the flanking zone of developing LRPs (Brenner et al., 2017) is also unique to this trajectory. Together, these data suggest that cells that were initially categorized as mature pericycle may be more accurately described as LRP-flanking pericycle cells, and this flanking fate is specifically induced by the initiation of a new root.

### Lateral root development shows strong signatures in endodermal transcriptomes

We next examined the impact of lateral root initiation on the endodermal cell file which is immediately exterior to the pericycle. We first re-clustered the 813 cell partition labeled as cortex and endodermis cells. This analysis revealed three distinct cell populations. The first population was a set of 127 cells which expressed cortex marker genes, the second population was a set of 410 cells which expressed endodermis marker genes, and the third population of 276 cells branching out from endodermis population (**Figure 5C, Supplemental Figure 10A, Supplemental Figure 10B**). This third population was transcriptionally most similar to the 410 cells expressing endodermis marker genes, and twenty-six percent of the cells in this third population were from the twenty hour timepoint. As a comparison, about eight percent of the 127 cell population were from the twenty hour timepoint, while about nine percent of the 410 cell population were from the twenty hour timepoint. This third population also had the highest expression of the NAC transcription factor, *ORESARA1* (*ORE1*) and autophagy marker gene *ATG8a* (**Figure 5D and E**). *ORE1* is a positive regulator of programmed cell death and autophagy marker gene ATG8a are specifically expressed in cells overlying LRPs (Escamez et al., 2020). *ORE1* expressing cells eventually die in order to make space for the lateral root to emerge (Escamez et al., 2020). As such, we categorized these three populations as cortex (127 cells), endodermis (410 cells), and lateral root-responding endodermis (LRE) (276 cells) (**Supplemental Table 1, Supplemental Figure 10A**).

To aid in identifying differentially expressed genes between endodermis and LRE, both cell types were compared to cortex cells as an outgroup. As in the XPP-LRP comparison, only genes that were called significantly different in at least two methods were called as DEGs (S**upplemental Figure 11**). This analysis yielded over 2000 DEGs between endodermis and cortex, and over 3000 DEGs between LRE and cortex. As expected, there was large overlap between these two sets of DEGs (**Supplemental Data 3**). A smaller set of genes were identified that were specific to the endodermis cells (204 genes), and specific to the LRE cells (303 genes), while 240 were commonly upregulated genes in endodermis and LRE (**Figure 5F**). The DEGs specific to the endodermis cells were enriched for GO terms associated with cell-cell junction assembly and water transport, while the DEGs specific to the LRE cells were enriched for GO terms associated with hormone response, auxin homeostasis, cell communication, lateral root development, and multiple stress responses (**Supplemental Data 3**).

Pseudotime analysis was performed to identify genes driving the transition from endodermis to the lateral root endodermis. This analysis showed a branch point between the two cell types. One branch that we termed the “main branch” was composed mostly of cells from the eight hour time point, and was transcriptionally similar to the rest of the endodermis cluster. The other branch that we called the “LRE branch” led towards the LRE cluster (**Supplemental Figure 12A**). Again, two separate trajectories were analyzed, one containing only endodermis cells and another containing endodermis cells below the branch point and LRE cells. This yielded a set of 2082 genes whose expression changed over the course of the cell developmental trajectory. After removing genes upregulated in cortex cells, a small set of genes were identified as specific to the main branch (154 genes) and to the LRE branch (648 genes) (**Supplemental Figure 12B, Supplemental Data 3**). GO analysis for the main branch specific genes showed enrichment for terms associated with responses to various stimuli. An example of a main branch specific gene is *DEEPER ROOT 1 (DRO1)* (**Supplemental Figure 12C**). Negatively regulated by auxin, *DRO1* is involved in gravity sensing in the root tip and determining lateral root branch angle (Uga et al., 2013; Guseman et al., 2017; Waite et al., 2020). *DRO1* loss of function mutants have increased horizontal lateral roots and have trouble establishing auxin gradients in response to gravistimulus. In our dataset, the majority of cells expressing *DRO1* are endodermal cells, suggesting these cells play a specific role in response to gravity. The LRE branch genes were enriched for GO terms associated with lateral root development, auxin homeostasis, auxin transport, and biosynthetic processes. Examples of LRE branch specific genes are *WRKY75* and *PINS-LIKE 5 (PILS5)* (**Supplemental Figure 12D**). *WRKY75* is induced particularly in phosphate starvation (Devaiah et al., 2007), and *PILS5* is an auxin efflux carrier regulating intracellular auxin homeostasis, both independently having a role in controlling root architecture. These along with gene ontologies pertaining to several biotic and abiotic stress responses (**Supplemental Data 3**) indicates that LRE cell population seems to be primed to dynamically assess environment and thereby regulate lateral root emergence.

## Discussion

Among the greatest mysteries in development is the process by which a stem cell begins proliferating and partitioning its progeny into increasingly determined cell fates. In plants, the initiation of lateral root development is among the best understood of these processes, yet many fundamental questions remain. Auxin is clearly a critical signal, but what other pathways interact with auxin response or regulate developmental steps downstream of auxin perception are still largely unknown. One key piece of missing information is a full accounting of transcriptional changes in the lateral root primordia during the critical window of initiation. In this study, we performed single-cell RNA sequencing at two timepoints on regions of roots where lateral root initiation was taking place. We identified cells of all cell types of the root outside the meristem in our population, including cells expressing lateral root markers. Through differential gene expression analysis, we identified genes upregulated in LRP cells as compared to XPP cells, many of which were indicative of cells undergoing cellular differentiation and organogenesis. We also identified a subset of pericycle and endodermal cells outside of the primordium itself that appear to be responding to the initiation of a new root.

We chose several genes for further study. Transcriptional reporters confirmed that most were expressed in early stage lateral roots, and that this expression was in many cases dependent on *ARF7* and *ARF19*. Several reporters showed additional expression in other parts of the root, including the primary root meristem. *OFP8* and *BRXL1* were expressed specifically in LRPs, making them potentially useful tools for studying and engineering root architecture. Repression of many of our candidate genes in XPP cells caused defects in lateral root development, a previously unknown role for some of them. For example, *HD2C* and *HD2B* have been linked to down-regulation of ribosomal biogenesis genes (Chen et al., 2018), a process we found to be strongly induced in lateral root primordia. This connection is consistent with our finding that repression of these genes led to higher density of lateral roots. It also highlights the fact that genes that are upregulated during primordia initiation may be important either for promoting developmental progression or for providing checks to keep cellular events coordinated or appropriately controlled. *OFP8* has been reported to act as a direct transcriptional repressor (Wang et al., 2011), and as a regulator of cell division patterns and organ shape via modification of the cytoskeleton (Snouffer et al., 2020).

Within the primordium itself, careful regulation of cell division and growth are essential for the formation of a primordium with normal morphology. The majority of our J0121^Col^≫dCas9R transgenic plants exhibited increased lateral root density or increased lateral root length as a proportion of total root length. These results suggest that these target genes may normally act as repressors of lateral root initiation or emergence. When they are repressed, development is accelerated. One interpretation is that cell division and lateral root development are the default state of competent pericycle cells. This would be analogous to the situation in the root epidermis where becoming a root hair is the default state that must be actively repressed in non-hair cells (Berger et al., 1998). This hypothesis is supported by experiments where laser ablation of surrounding tissue causes unrestricted cell division in the pericycle cell file (Marhavý et al., 2016), and where exogenous auxin treatments trigger lateral root initiation in every pericycle cell (Himanen et al., 2002).

Using cell-type specific repression of our candidates in XPP cells allowed us to avoid several problems with global mutant analyses. For example, *PLT3* and *BRXL1* play critical roles in the primary root meristem, making interpretation of any lateral root phenotypes difficult. A narrower scope of repression may also reduce the likelihood of compensation from paralogs. The repression lines decrease expression only in XPP cells and in LRP cells up to stage IV. This tissue-specific expression may explain the phenotypic difference between our *PLT3*-repressed line and *plt3 plt5 plt7* mutants, which show decreased lateral root emergence (Du and Scheres, 2017). It is possible that *PLT3* is playing opposing roles in different cell files of the root or at different points in lateral root development, depending on the interacting partners that are present in these cell files at these developmental stages. Alternatively, the phenotypes we observed could reflect feedback effects on other genes in the same gene regulatory network. The analysis performed here is likely to have missed phenotypes, especially those that rely on complex interactions with a soil environment or on a different set of metrics (Fitter, 1987; Lynch, 1995). For example, the irregular spacing and rate of growth of lateral roots in our BRXL1/BRXL4-repressed lines did not significantly impact the metrics we tested (**Supplemental Figure 9**). Introgression of J0121 into different accessions would allow access to a broader array of cryptotypes (Chitwood and Topp, 2015; Ristova et al., 2018), and a more holistic view of impacts on root architecture.

We found that our single-cell experiments captured the majority of previously reported LRP-enriched transcripts in our LRP-assigned cells, and these transcripts were not enriched in our XPP-assigned cells. We additionally found many more LRP-enriched genes in our library than in bulk transcriptomes, underlining the utility of this method in examining rare developmental events. Many of our XPP-upregulated genes oscillate antiphase to auxin response in the basal meristem. These antiphase-oscillatory genes are upregulated in cells that are *not* competent to form lateral roots. They may be actively preventing lateral root initiation in XPP cells, in alignment with the hypothesis that LRP-competency may be the default state of XPP cells. Notably, there are also 698 cells within our stele cluster that were too ambiguous in their gene expression to assign a cell label. It is possible some of these cells are precursors to stele cells, such as XPP precursors that are not yet lateral root competent. How distinct lateral root competent and non-competent XPP cells are is unknown—the only known distinguishing feature between the two is a characteristic auxin response maximum or lack thereof. Additional differential gene expression between different timepoints within the XPP cluster and this non-assigned stele cell cluster may yield novel insights.

Though lateral root development is specific to xylem pole pericycle cells, the process is not cell-autonomous. Our analysis identified a population of endodermal cells distinct from the main endodermis branch. These cells were enriched in the expression of genes falling in ontology categories for hormone and auxin response, cell-cell communication, and lateral root development, making a strong case that they are responding to developing primordia in underlying pericycle cells. They were also enriched in *ORE1* expression, a gene that has recently been shown to play a role in lateral root initiation and emergence through programmed cell death of tissue overlying LRPs (Escamez et al., 2020). Consequently, this population of endodermal cells appears to be responding to lateral root initiation in neighboring pericycle cells, and forging a path for the incipient primordium.

We were also able to identify a subset of pericycle cells that were likely directly adjacent to the primordium and responding with a distinct transcriptional program that included a combination of auxin-repressing and cytokinin-induced genes. A dynamic analysis of the gene regulatory network governing lateral root development established that early cell fate determining genes initiate multiple genetic feedback loops that divide the developing primordium into two zones, a central proliferative core and flanking cells that have inhibited expression of meristematic genes to repress cell division (Lavenus et al., 2015). Notably, PPP cells in our dataset also show strong expression of several auxin-inhibitory genes. PPP cells never initiate lateral root development, even though these cells receive the same auxin maximum signal as XPP cells during specification. It is possible these repressors of auxin signaling act to prevent spurious root development in multiple cell files.

As human activity changes the climate and environments in which plants grow, understanding root development will help us engineer crops that are more robust to nutrient scarcities and environmental extremes. The major pathway by which eudicot plants regulate their root architecture is through modification of the position, spacing, density, and length of lateral roots. The stages of lateral root development are regulated by distinct genetic circuitry. Every stage represents an opportunity for natural and engineered modification of this developmental process. Molecular characterization of early stages of lateral root at single-cell resolution gives us a more comprehensive understanding of this fate decision and the molecular pathways that tune it.

---

**Supplemental Figure 1. Marker gene expression profiles and stele cell UMAP.** A. Heatmap (column-scaled) visualizing average normalized expression of marker genes in the columella cells, epidermis cells, cortex & endodermis cells, and stele cells. Scale bar represents the z-score of the normalized expression values. B. Heatmap (column-scaled) visualizing average normalized expression of marker genes in different stele cell types. Scale bar represents the z-score of the normalized expression values. C. UMAP of stele cells colored by experiment. D. Fraction of xylem pole pericycle (XPP), lateral root primordia (LRP), mature pericycle (MP), and phloem pole pericycle (PPP) cells from each experiment.

**Supplemental Figure 2. Xylem pole pericycle developmental trajectories.** A. UMAP of the XPP to Mature Pericycle trajectory colored by pseudotime. B. UMAP of the XPP to LRP trajectory colored by pseudotime. C. Expression UMAP (XPP and Mature Pericycle cells) of DEGs identified in both trajectories (XPP to Mature Pericycle and XPP to LRP) and in only the XPP to Mature Pericycle trajectory. D. Expression UMAP (XPP and LRP cells) of DEGs identified in both trajectories and in only the XPP to LRP trajectory. E.. Heatmap (row-scaled) visualizing average normalized expression of genes identified as differentially expressed in the XPP to LRP trajectory. Scale bar represents the z-score of the normalized expression values.

**Supplemental Figure 3. Size-adjusted Venn diagram visualizing overlap of genes between different DEG calling methods.** For each circle of the Venn Diagram, the total number of DEGs as well as the number of DEGs up in LRP and up in XPP are shown. All genes that were called in two or more methods were used for downstream analysis.

**Supplemental Figure 4. Comparison of XPP and LRP DEGs from the single-cell library to bulk transcriptomes.** A. Comparison to LRP-induced genes from Vanneste *et al*, 2005. B. Comparison to time-course analysis of lateral root initiation at nine and twenty-one hours post-bend from Voß *et al*, 2015. C. Comparison to auxin-induced and auxin-repressed genes in the root from Lewis *et al,* 2013. D. Comparison to ARF19-specific auxin-induced genes from Powers *et al*, 2019. E. Comparison to genes oscillating in phase and antiphase to auxin in the basal meristem during lateral root specification from Moreno-Risueno *et al*, 2010. F. Comparison to auxin-induced genes during early lateral root development from Ramakrishna *et al,* 2019. Each gene set from bulk transcriptomes is compared to the XPP and LRP DEGs with a size-adjusted Venn diagram. The number of genes in each mutually-exclusive area of each Venn diagram are specified. The values in purple and turquoise denote the hypergeometric distribution for the union XPP/bulk transcriptome and LRP/bulk transcriptome respectively.

**Supplemental Figure 5. Design of J0121^Col^≫dCas9R system to generate cell type-specific dCas9-repressor mediated knockdown of candidate gene expression.** A. J0121^Col^ is an enhancer trap line where the UAS-Gal4 system drives expression of GFP in the xylem pole pericycle cell file, visualized in confocal microscopy image and B. labeled in cartoons (in green). C. Top panel indicates the enhancer trap cassette in the J0121^Col^ and bottom panels is design of perturbation plasmids included up to three guide RNAs, tagged as location L1, L2, and L3, and a UAS promoter driving expression of dCas9-repressor cassette. Perturbation plasmids with respective cloned guide RNAs were transformed into J0121^Col^ background to drive repression of target genes specifically in xylem pole pericycle cells.

**Supplemental Figure 6. Example seedling traces from J0121^Col^≫dCas9R perturbation lines.** Roots from T2 perturbation line seedlings were quantified for various lateral root developmental phenotypes using SmartRoot.

**Supplemental Figure 7. Transcriptional reporters of chromatin regulator candidate genes in wild type and** *arf7arf19* **roots.** Fluorescent microscopy images of transgenic plant lines carrying transcriptional reporters of candidate genes in early stage lateral root primordia are shown in the large left-side image of each panel. Smaller images of the same transcriptional reporters in a region of the differentiated zone of the root without any developing primordia (above) and in the root apical meristem (below) are shown on the right in each panel. The same reporters in the same regions of the root are imaged in *arf7arf19* mutant background plants on the right. The number of independent transgenic lines imaged per construct and the number of plants within each line that showed expression are reported at the bottom. The lower panel represents 1000 bp upstream of the transcription start site for each gene, with auxin response elements (TGTC/GACA) highlighted in red. Yellow bars indicate CDSs from other genes. This panel was obtained from http://bar.utoronto.ca/cistome.

**Supplemental Figure 8. Transcriptional reporters of cell cycle candidate genes in wild type and** *arf7arf19* **roots.** Fluorescent microscopy images of transgenic plant lines carrying transcriptional reporters of candidate genes in early stage lateral root primordia are shown in the large left-side image of each panel. Smaller images of the same transcriptional reporters in a region of the differentiated zone of the root without any developing primordia (above) and in the root apical meristem (below) are shown on the right in each panel. The same reporters in the same regions of the root are imaged in *arf7arf19* mutant background plants on the right. The number of independent transgenic lines imaged per construct and the number of plants within each line that showed expression are reported at the bottom. The lower panel represents 1000 bp upstream of the transcription start site for each gene, with auxin response elements (TGTC/GACA) highlighted in red. Yellow bars indicate CDSs from other genes. This panel was obtained from http://bar.utoronto.ca/cistome.

**Supplemental Figure 9. Transcriptional reporters of stemness candidate genes in wild type and** *arf7arf19* **roots.** Fluorescent microscopy images of transgenic plant lines carrying transcriptional reporters of candidate genes in early stage lateral root primordia are shown in the large left-side image of each panel. Smaller images of the same transcriptional reporters in a region of the differentiated zone of the root without any developing primordia (above) and in the root apical meristem (below) are shown on the right in each panel. The same reporters in the same regions of the root are imaged in *arf7arf19* mutant background plants on the right. The number of independent transgenic lines imaged per construct and the number of plants within each line that showed expression are reported at the bottom. The lower panel represents 1000 bp upstream of the transcription start site for each gene, with auxin response elements (TGTC/GACA) highlighted in red. Yellow bars indicate CDSs from other genes. This panel was obtained from http://bar.utoronto.ca/cistome.

**Supplemental Figure 10. Marker gene expression profiles and experiment breakdown of cortex, endodermis, and lateral root endodermis cells.** A. Heatmap (column-scaled) visualizing average normalized expression of marker genes in the cortex, endodermis, lateral root endodermis (LRE) cells. B. Fraction of cortex, endodermis, and LRE cells from each experiment.

**Supplementary Figure 11. DEG overlaps with different methods for Endodermis/Lateral Root Endodermis analyses** A. Endodermis vs cortex comparison and B. LRE vs cortex comparison. For each ensemble of the Venn Diagram, the total number of DEGs, the number of DEGs up in Cortex and up in Endodermis (for A) and LRE (for B) are added to the diagram.

**Supplemental Figure 12. Pseudotime analysis of Endodermis to Lateral Root Endodermis Cells**. A. UMAP of the endodermis to LRE trajectory colored by pseudotime. B. UMAP of the expression of gene sets that differed significantly as a function of pseudotime in the main endodermis branch and the LRE Branch. C. Expression UMAP of DRO1. D. Expression UMAPs of WRKY75 and PILS5.

**Supplementary Table 1. Breakdown of cell types by experiment.**

## Author contributions

A.L., H.P.G., C.Q., J.T.C and J.L.N. designed the research; H.P.G., A.L., S.G., J.C.C., J.E.Z., W.G. and J.T.C performed research; A.L., H.P.G., K. J-B., J.T.C and J.L.N. analyzed data; and A.L., H.P.G., K. J-B. and J.L.N. wrote the manuscript.

## Acknowledgements

We would like to thank members of the Nemhauser, Queitsch, Trapnell, Steinbrenner and Imaizumi groups for helpful discussions and technical guidance. We are grateful for the generosity of Dr. Dolf Weijers and his group in hosting A.L. to work on TMO6, as well as for Dr. Bert De Rybel’s sharing of TMO6 reporter lines. This work was supported by the National Institutes of Health (R01-GM107084; J.L.N.; R01-GM079712, C.Q. and J.C.), the Howard Hughes Medical Institute Faculty Scholars Program (J.L.N.), and the National Science Foundation (IOS-1748843; C.Q. and J.C.). A.L. was supported by an NSF Graduate Research Fellowship (DGE-1256082), and S.G. was supported by an EMBO Postdoctoral Award (ALTF 409-2019).

## Methods

### Construction of plasmids

Each reporter plasmid is composed of the selected promoter, the red fluorescent protein mScarlet with a nuclear localization tag (Bindels et al., 2017), and the rbcS terminator (Siligato et al., 2016). The three parts were assembled using golden gate assembly in the modified pGII-Hygr vector containing compatible Golden Gate sites (Weber et al., 2011). For each of the ten constructed reporters, the promoter sequence of the reporter corresponds to the DNA sequence in 5’ of the start codon of the corresponding gene based on TAIR10 genome from http://plants.ensembl.org/. While aiming for a 2000 bp length, the lengths of promoters are usually smaller to avoid coding sequences of other genes. The exact sequence and length of the selected promoter for each of the 10 genes can be found in supplementary material (**Supplementary Data 2**). The promoter sequence for the TMO6 reporters corresponds to the sequence used in a previous work from (Smet et al., 2019). The promoter sequences were amplified from purified *Arabidopsis thaliana* Col-0 genomic DNA using Q5 polymerase and with primer adding the specific golden gate spacer. After gel purification, each promoter part was cloned and sequence verified in a pBLUNT entry vector. Three part Golden Gate assembly was performed using the pBLUNT promoter plasmid, mScarlet, rbcS terminator to clone reporter plasmid.

For cell type specific knockdown mediated by J0121^Col^≫dCas9R, Gibson cloning was used to replace the egg specific promoter and Cas9 from pHEE401E (Wang et al., 2015) with UAS promoter and dCas9-TPL fusion (Khakhar et al., 2018). The resulting plasmid is used as starting point to clone two or three guide RNA against the promoters of selected gene/genes (identified using CHOP CHOP (Montague et al., 2014) ranging from −200 to +100 region from the annotated TSS) using PCR and golden gate strategy described in (Wang et al., 2015).

### Plant growth conditions and sample preparations

For all plant experiments, *Arabidopsis* seeds were sown on 0.5× LS 0.8% agar plates, stratified at 4°C for 2 days, and grown in continuous light conditions at 22°C for respective experimental design.

### Microdissection of root bend and protoplast isolation

For lateral root induction assays, ~150 seedling for each timepoint and treatment were rotated 90° 4 days post-germination (dpg) or in the case of the control treatment, the primary root tip was marked at this time and the plates were not turned. On the day of single-cell library preparation first the protoplasting enzyme mix was prepared adapted from (Yoo et al., 2007). Briefly, 20 mM MES (pH 5.7) containing 1.25% (wt/vol) cellulase R10 (C224 PhytoTechnology Laboratories), 0.3% (wt/vol) macerozyme R10 (M481 PhytoTechnology Laboratories,), 0.4 M mannitol and 20 mM KCl was prepared and incubated in 55°C warm water bath for 10 minutes. Upon cooling to room temperature (~25 °C),10 mM CaCl2, 1–5 and 0.1% BSA was added. Root bends (or marked region in no bend control) were microdissected using a scalpel at eight hours (control and eight hour treatment groups) and 20 hours (20 hour treatment group) post-bending, approximately 1 mm from the bend or mark in both directions. Using fine forceps dissected bend tissue was transferred into 30 mm dishes containing 1 mL of protoplasting enzyme mix and gently scored using a fresh scalpel to increase exposure of interior cell files to protoplasting enzymes. The plates were then flooded with 9 mL more protoplasting enzyme mix and incubated at room temperature for one hour with gentle shaking (75-80rpm). Protoplasting enzyme mix was filtered through 40 μm cell strainer, transferred and centrifuged in 15 ml conical tubes for five min at 500g. The supernatant was carefully removed and resuspended in 50 μl protoplasting mix without enzymes. Cell number was determined by hemocytometer and density was adjusted to ~1000cell/μL

### Construction and selection of transgenic Arabidopsis thaliana lines

Floral dip (Clough and Bent, 1998) was used to introduce constructs into Col-0 and *arf7arf19* lines (Okushima et al., 2007). T1 seedlings were selected on 0.5X LS (Caisson Laboratories, Smithfield, UT) + 25μg/ml Hygromycin B + 0.8% bacto-agar. Plates were stratified for two days, exposed to the light for six hours, and then grown in the dark for three days (Harrison et al., 2006). Hygromycin resistant seedlings were identified by their long hypocotyl, enlarged green leaves, and long root. Transformants were transferred on soil, and T2 seeds were collected.

### Lateral root bend assay and confocal microscopy

For each reporter, one Col-0 T1 line representative of other characterized T1 lines was selected to perform lateral root bend essay. For each reporter, 20 T2 seeds of the corresponding T1 line were placed on 0.5 LS+0.8% bacto-agar plate following a specific pattern to avoid seedling collision during the lateral root bend essay. The plate was stratified during 120 hours, grown vertically for 96 hours at 22°C, rotated 90°C while keeping vertically and grown for an additional 20 hours.

### Confocal microscopy of reporter lines at root bends

Seedlings were fixed at 4 dpg + 20 hours using 4% formaldehyde using vacuum infiltration followed by cleared using ClearSee solution (Kurihara et al., 2015). Fixed and cleared seedlings were mounted on microscopic slides using 50% glycerol and parafilm edges to avoid coverslips pressing on the root. Seedlings were imaged at the bend region using a SP5 confocal microscope. Images were processed using FIJI.

### Comparison between Col-0 and arf7arf19 lines

To perform comparative imaging of Col-0 and *arf7arf19* reporter lines, seeds of selected T1 lines for both Col-0 and *arf7arf19* reporter lines were placed on the same 0.5 LS+0.8 phytoagar plate. The selected Col-0 and *arf7arf19* lines for each reporter is specified below the microscope images of supplemental figure 7 to 9 as being highlighted in bold. Plates were stratified for 2 days, and grown vertically at 22°C for 10 days. Then, seedlings were imaged using a Leica DMI 3000B microscope at the root tip region, at the region above the root tip corresponding to the initiation of root hair and at the lateral root initiation region. As the *arf7arf19* line does not develop lateral roots, the theoretical lateral root initiation region is identified by identifying a lateral root primordium in the Col-0 seedling and imaging at a similar region.

### Lateral root phenotypes of repression lines

For cell type specific knockdown mediated by J0121^Col^≫dCas9R we leveraged an established GAL4-UAS system (Laplaze et al., 2005) of enhancer trap line J0121. We backcrossed the J0121 line, made in the C24 background, eight times into the Col-0 background to produce a strain we refer to as J0121^Col^. We confirmed that J0121^Col^ retained strong GFP expression in xylem pole pericycle and exhibited Col-0-like root growth dynamics. Transformants were selected as described above and T2 seeds for at least 10 lines were collected. T2 seeds were grown vertically for 10dpg in 100mm square plate on 0.8% bacto agar and were scanned on a flatbed scanner (Epson America, Long Beach, CA) for phenotyping. Since the T2 generation is a segregating population for the transformed plasmid, seedlings were genotyped for the presence of vector backbone to identify positive seeding. Roots for positive seedlings were traced using ImageJ and SmartRoot plugin (Lobet et al., 2011) and analyzed and plotted using R package archiDART (Delory et al., 2016) package plot were generated using ggplot2. Density was measured as the total number of lateral roots divided by the length of the primary root. Proportion of lateral root length was measured as the summed length of all lateral roots divided by the summed length of all lateral roots and the primary root length.

### Single-cell RNA-sequencing Protocol

Single-cell RNA-Seq was performed using the 10X scRNA-Seq platform, the Chromium Single Cell Gene Expression Solution (10X Genomics). Two replicates were produced for each timepoint of the experiment for a total of six samples. We also generated two replicates from a transgenic plant line that slows the rate of degradation of IAA14 (Guseman et al., 2015), dissecting root bends in this line twenty hours after bending. This line shows delayed lateral root development, and we initially thought to compare its transcriptomes to our wild type treatment groups. Unfortunately, one of the replicates of this line failed at the 10X droplet-binding stage, so we did not obtain the same number cells from this treatment group as from our other groups. Consequently, we excluded these cells from further analysis.

### Estimating Gene Expression in Individual Cells

Single-cell RNA-sequencing reads were sequenced using an Illumina NextSeq 500 and then mapped to the TAIR10 *Arabidopsis* genome using the software Cellranger (v.3.0.1). Cellranger produces a matrix of UMI counts where each row is a gene and each column represents a cell. The ARAPORT gene annotation was used. For the analysis, reads from two 00 hour replicates, two 08 hour replicates, and two 20 hour replicates were aggregated using the aggr command in cellranger to normalize to an equivalent number of mean reads per cell across samples. This resulted in a mean of 14,516 reads per cell, a median of 1,411 genes per cell, and a median of 2,873 UMIs per cell.

### Running Monocle 3: Dimensionality Reduction, and Cell Clustering

The filtered output of the Cellranger pipeline (../outs/filtered_gene_bc_matrices_mex/) was parsed into R (v. 3.5.0). Particularly the matrix.mtx file was parsed using the readMM() function from the Matrix package (https://cran.r-project.org/web/packages/Matrix/Matrix.pdf), and the barcodes.tsv file and the genes.tsv file were parsed using the read.table() function. Genes that were expressed in less than 10 cells were removed from the analysis. In addition, the 346 genes induced due to protoplast generation process were also removed from the analysis (Birnbaum et al., 2003). The barcodes table was updated to label cells by Sample Number and Experiment. Finally the expression matrix, the barcode table, and the gene table were converted into a CellDataSet (CDS) using the new_cell_data_set() function in Monocle 3 (cole-trapnell-lab/monocle3, 2020) (v. 0.1.2; https://cole-trapnell-lab.github.io/monocle3/)

All Monocle 3 analysis was performed on a High Performance Computing cluster using 128 GB of RAM spread across eight cores. We visualized cell clusters and trajectories using the standard Monocle workflow. Monocle internally handles all normalization needed for dimensionality reduction, visualization, and differential expression. The CDS was normalized and pre-processed using the preprocess_cds() function with the following parameters:

~~~
num_dim=100,
method=“PCA”,
norm_method=“log”,
scaling=T,
residual_model_formula_str=“~ Sample_Number”
~~~

Preprocessing involves reducing the dimensionality of the data (the number of genes) using principal component analysis (PCA). Here, we retain the first 100 PCs for further dimensionality reduction, in addition we reduce batch effect across samples. Then, the PCA matrix was used to initialize a nonlinear manifold learning algorithm implemented in Monocle 3 called Uniform Manifold Approximation and Projection ( UMAP) (McInnes et al., 2018). This allows us to visualize the data into two or three dimensions. Specifically, we projected the data onto two dimensions using the reduce_dimension() function using the following parameters:

~~~
reduction_method=“UMAP”,
preprocess_method=“PCA”,
umap.metric=“cosine”,
umap.min_dist=0.1,
umap.n_neighbors=15L,
umap.nn_method=“annoy”,
umap.fast_sgd=F
~~~

Next, cells were clustered into partitions using PhenoGraph (Levine et al., 2015) and into clusters using the Leidan community detection algorithm (Traag et al., 2019) with the cluster_cells() function using the following parameters:

~~~
reduction=“UMAP”,
k=20,
louvain_iter=1,
partition_qval=0.05,
weight=T,
resolution=c(10^seq(−6,0)).
~~~

This resulted in 5 cell partitions, and 43 cell clusters. Next, for each cell partition, a cell trajectory was drawn atop the projection using Monocle’s reversed graph embedding algorithm, which is derived from SimplePPT (Mao et al., 2017) using the learn_graph() function using the following parameters:

~~~
use_partition=T
close_loop=F,
learn_graph_control=list(prune_graph=T)
~~~

To further analyze the partition we annotated as stele, those cells were reclustered together and were reanalyzed using Monocle 3 as previously described except during preprocessing, instead of performing PCA on all the genes, PCA was performed on just a set of stele cell marker genes reported earlier (Mao et al., 2017). Specifically the “use_genes” option was used in the preprocess_cds() function, and a list of gene names was provided. Lastly, cells were clustered using the cluster_cells() function, and the “resolution” parameter was set to 0.001. This produced 3 partitions, and 7 clusters.

To further analyze the clusters we annotated as Phloem Pole Pericycle, Mature Pericycle, Xylem Pole Pericycle, and Lateral Root Primordia, those cells were reclustered together and were reanalyzed using Monocle 3 as previously described except during preprocessing, instead of performing PCA on all the genes, PCA was performed on just a set of stele cell marker genes reported in (Brady et al., 2007). Again, Specifically the “use_genes” option was used in the preprocess_cds() function, and a list of gene names was provided. Next, the data was reduced onto two dimensions using the reduce dimension function but the “umap.min_dist” parameter was set to 0.01. Afterwards cell clusters were called as before using the cluster_cells() function except the “resolution” parameter was set to 0.0005. Next a cell trajectory was created using the learn_graph() function. Finally cell clusters were recalled using the cluster_cells() function except the “resolution” parameter was set to 0.001. This produced 1 partition and 4 clusters. To further analyze the partition we annotated as Cortex and Endodermis, those cells were reclustered together and were reanalyzed using Monocle 3 as previously described except during preprocessing, instead of performing PCA on all the genes, PCA was performed on just a set of cortex and endodermis cell marker genes reported in (Brady et al., 2007). This produced 2 partitions and 15 clusters.

### Estimating Doublets

Single Cell Remover of Doublets (Scrublet) was used to predict doublets in our scRNA-Seq data (Wolock et al., 2019). Using python 3.5, Scrublet was run using default settings as described by the example tutorial that is available as a Python notebook (https://github.com/AllonKleinLab/scrublet/blob/master/examples/scrublet_basics.ipynb). The only significant change was that expected double rate was set to 0.1; in the tutorial it is 0.06.

### Assigning Cell Types

A set of known marker genes derived from earlier studies using green fluorescent protein (GFP) marker lines of the *Arabidopsis* root were used to identify cell types (Brady et al., 2007; Cartwright et al., 2009). The average gene expression of each marker set was used to assign cell types to cells, with cells being assigned the cell type it had the highest average expression

### Calling Differentially Expressed Genes: Xylem Pole Pericycle vs. Lateral Root Primordia

Differentially expressed genes between the cluster of cells labeled Xylem Pole Pericycle and the cluster of cells labeled Lateral Root Primordia were called using three different approaches.

The first approach involved running a generalized linear model to predict the average log express of each gene as a function of the cell type label. This was done using a subsetted CDS containing only the Xylem Pole Pericycle cells and the Lateral Root Primordia cells, and the Monocle 3 function fit_models() with the following parameters:

~~~
model_formula_str = “~cell_type”,
expression_family=“negbinomial”,
clean_model=T
~~~

where “cell_type” is a column in the dataframe returned by the colData() function that describes the cell type label associated with a cell/barcode. Using an FDR cutoff of 0.1, 1204 genes were called as differentially expressed between Xylem Pole Pericycle and Lateral Root Primordia. Of these, 424 were more highly expressed in Xylem Pole Pericycle, and 780 were more highly expressed in Lateral Root Primordia.

The second approach involved using the Mann-Whitney-Wilcoxon test to determine if the rank-sum of the normalized expression values for each gene differed between the Xylem Pole Pericycle population and the Lateral Root Primordia population. Mann-Whitney-Wicoxon test p-values were adjusted for multiple test comparisons using the Benjamini-Hochberg procedure via the R function p.adjust() from the stats package. Normalized expression values were calculated by taking the UMI matrix, obtained using Monocle 3’s counts() function and dividing by the size factors of each cell using Monocle 3’s size_factors() function. Using an adjusted p-value cutoff of 0.0001, 2088 genes were called as differentially expressed with 604 genes more highly expressed in Xylem Pole Pericycle and 1484 genes more highly expressed in Lateral Root Primordia.

The last approach involved using the analysis tool Vision. The normalised expression matrix for only XPP and LRP cell clusters was exported from Monocle. The gene signature of pericycle cell population from (Parizot et al., 2012) were used for running Vision() and analyse() function. LRP cell population was selected in browser view mode to identify DEG against XPP population. Vision identified 4900 DEGs using an FDR of less than 0.05.

### Pseudotime Analysis: Xylem Pole Pericycle Cells Development

Pseudotime analysis was performed on two subsetted CDSs, one containing only Xylem Pole Pericycle cells and Mature Pericycle cells, and the other containing only Xylem Pole Pericycle cells and Lateral Root Primordia cells. Cells in both CDSs were assigned a pseudotime on the cell trajectory using Monocle 3’s order_cells() function with the Xylem Pole Pericycle serving as the root of the trajectory. Genes whose expression changed as a function of pseudotime were identified using a generalized linear model. This was done on both CDSs using the fit_models() function and the following parameters:

~~~
model_formula_str = “~pseudotime”,
expression_family=“negbinomial”,
clean_model=T
~~~

Using an FDR cutoff of 0.1, 1394 genes were identified as changing as a function of pseudotime in the CDS containing only Xylem Pole Pericycle cells and Mature Pericycle cells, and 1014 genes were identified as changing as a function of pseudotime in the CDS containing only Xylem Pole Pericycle cells and Lateral Root Primordia cells with an overlap of 510 genes.

### Calling Differential Expressed Genes: Endodermis vs. Lateral Root Primodia Responding Endodermis

As previously described, a generalized linear model was used to identify differentially expressed genes between the cluster of cells labeled Endodermis, and the cluster of cells labeled Lateral Root Primodia Responding Endodermis. Using an FDR cutoff of 0.1, 1251 genes were identified as differentially expressed, with 748 genes being more expressed in Endodermis and 503 genes being more expressed in Lateral Root Endodermis. To identify additional DEGs, the MMW test was performed comparing Endodermis to Cortex and Lateral Root Endodermis to Cortex.

### Pseudotime Analysis: Endodermis Cells

Pseudotime analysis was performed on two subsetted CDSs, one with only Endodermis cells, and the other containing only Endodermis below the branch point, and Lateral Root Primodia Responding Endodermis. As previously described, cells were assigned a pseudotime along the cell trajectory with the Endodermis cells below the branch point serving as the root. As previously described, a generalized linear model was used to identify differentially expressed genes as a function of pseudotime. Using an FDR cutoff of 0.1, 2063 genes were identified in the CDS with only Endodermis, and 2079 genes were identified in the CDS with only Endodermis and LRP Responding Endodermis with an overlap of 2060 genes.

### GO Term Enrichment Analysis

GO term enrichment analysis was performed using PANTHER (http://pantherdb.org/) (Mi et al., 2019). For GO term enrichments for XPP, LRP, Endodermis, and LRE, genes that were significant in at least 2 methods were used for analysis. All genes in the *Arabidopsis* genome were used as a background. Fisher’s Exact test was used and the False Discovery Rate was calculated for multiple test correction. The complete annotation data set for biological process, molecular function, and cellular component GO terms were used for analysis.

### Accession Numbers

The GEO accession number for the scRNA-seq data reported in this paper is GSE158761 (https://www.ncbi.nlm.nih.gov/geo/query/acc.cgi?acc=GSE158761).

## Supplementary Data files

Supplemental Data 1. Xylem Pole Pericycle, Lateral Root Primordia, and Mature Pericycle DEG analysis sheet and GO Terms

Supplemental Data 2. Cloning Primers for generation of plasmids used to generate reporter and J0121^Col^≫dCas9R transgenic lines

Supplemental Data 3. Cortex, Endodermis, and Lateral Root Endodermis DEG analysis sheet and GO Terms

