## Supplemental Figures for "A single cell view of the transcriptome during lateral root initiation in *Arabidopsis thaliana*"

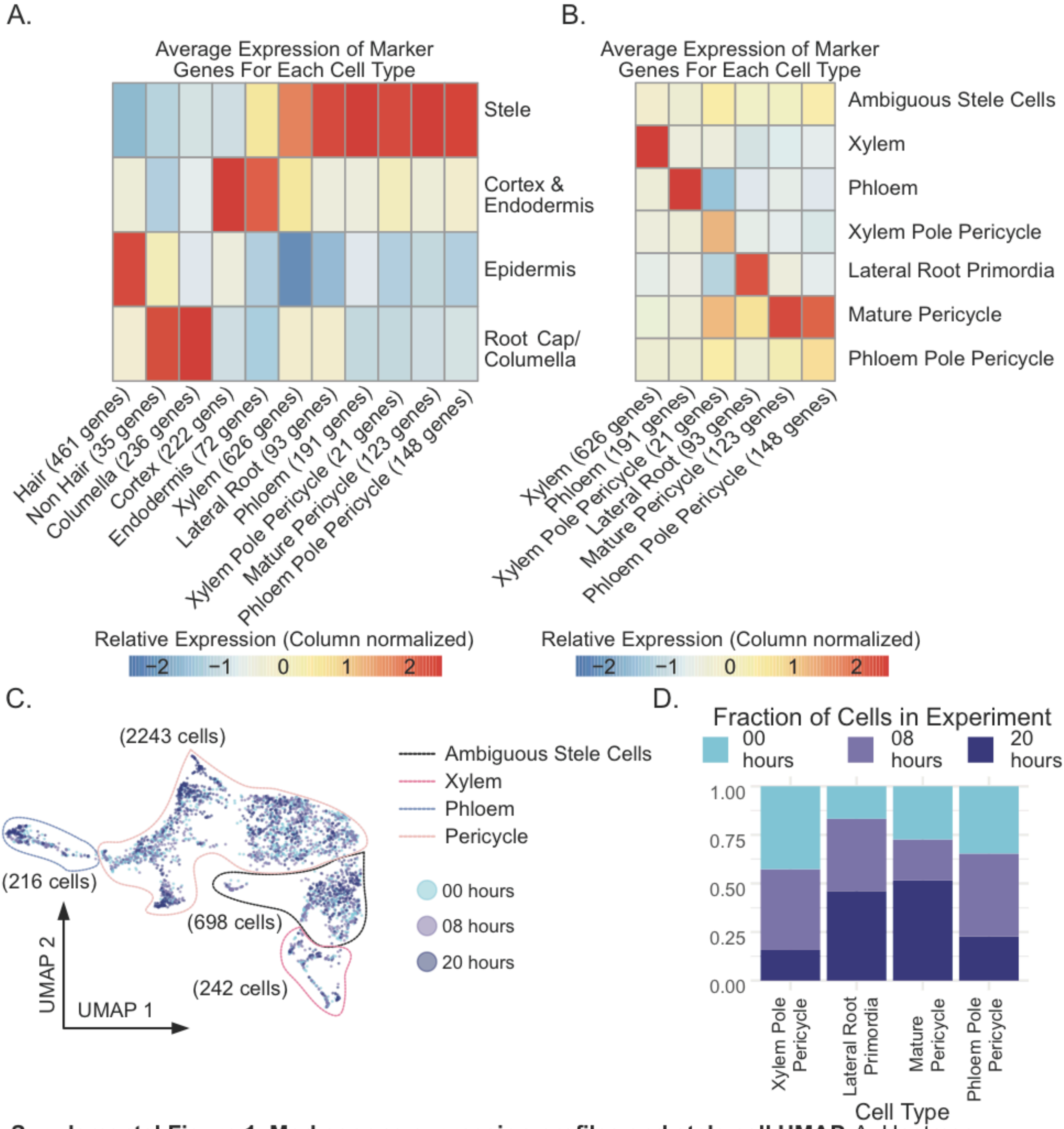


**Supplemental Figure 1. Marker gene expression profiles and stele cell UMAP.** A. Heatmap (column-scaled) visualizing average normalized expression of marker genes in the columella cells, epidermis cells, cortex & endodermis cells, and stele cells. Scale bar represents the z-score of the normalized expression values. B. Heatmap (column-scaled) visualizing average normalized expression of marker genes in different stele cell types. Scale bar represents the z-score of the normalized expression values. C. UMAP of stele cells colored by experiment. D. Fraction of xylem pole pericycle (XPP), lateral root primordia (LRP), mature pericycle (MP), and phloem pole pericycle (PPP) cells from each experiment.

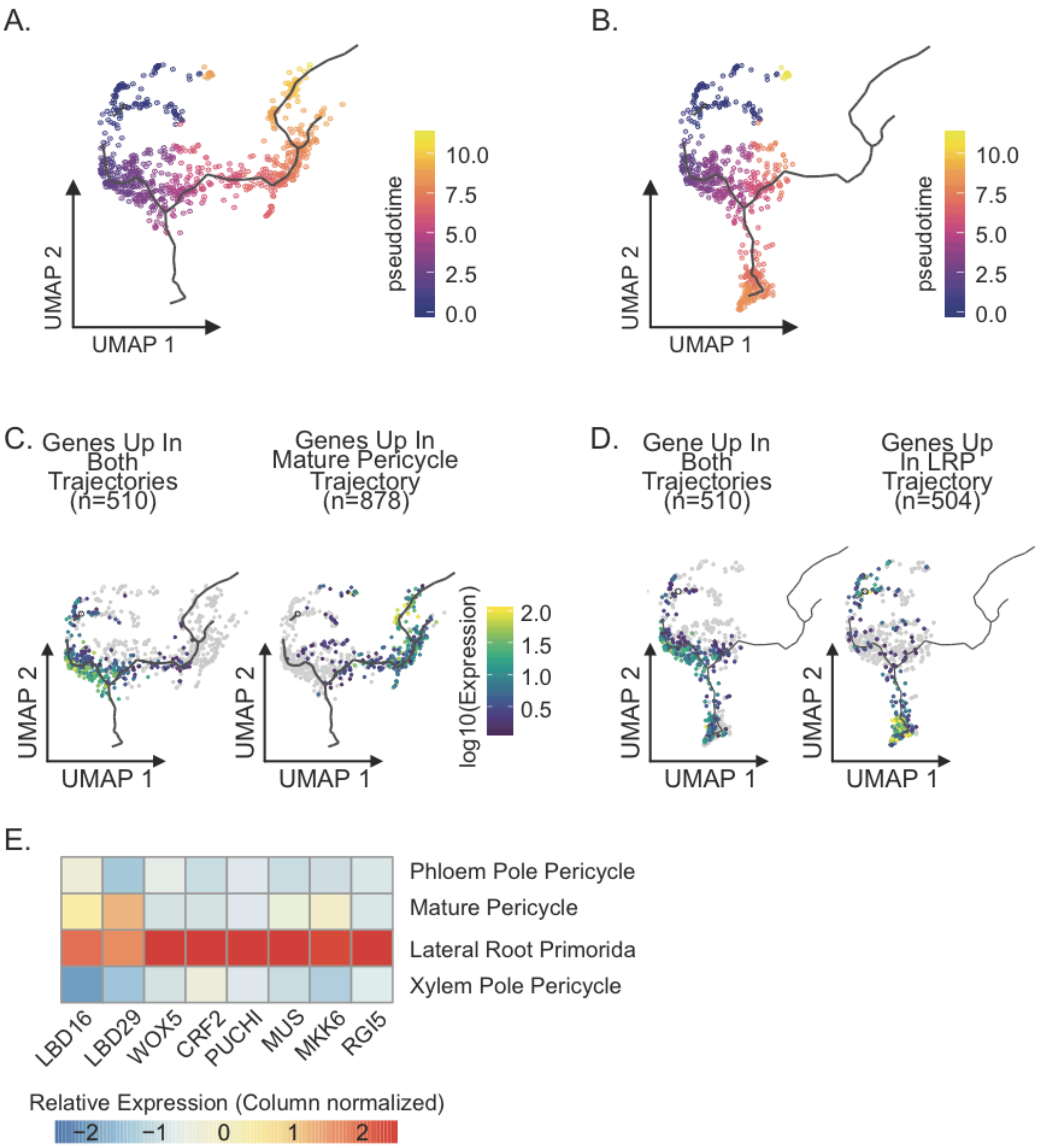


**Supplemental Figure 2. Xylem pole pericycle developmental trajectories.** A. UMAP of the XPP to Mature Pericycle trajectory colored by pseudotime. B. UMAP of the XPP to LRP trajectory colored by pseudotime. C. Expression UMAP (XPP and Mature Pericycle cells) of DEGs identified in both trajectories (XPP to Mature Pericycle and XPP to LRP) and in only the XPP to Mature Pericycle trajectory. D. Expression UMAP (XPP and LRP cells) of DEGs identified in both trajectories and in only the XPP to LRP trajectory. E. Heatmap (column-scaled) visualizing average normalized expression of genes identified as differentially expressed in the XPP to LRP trajectory. Scale bar represents the z-score of the normalized expression values.

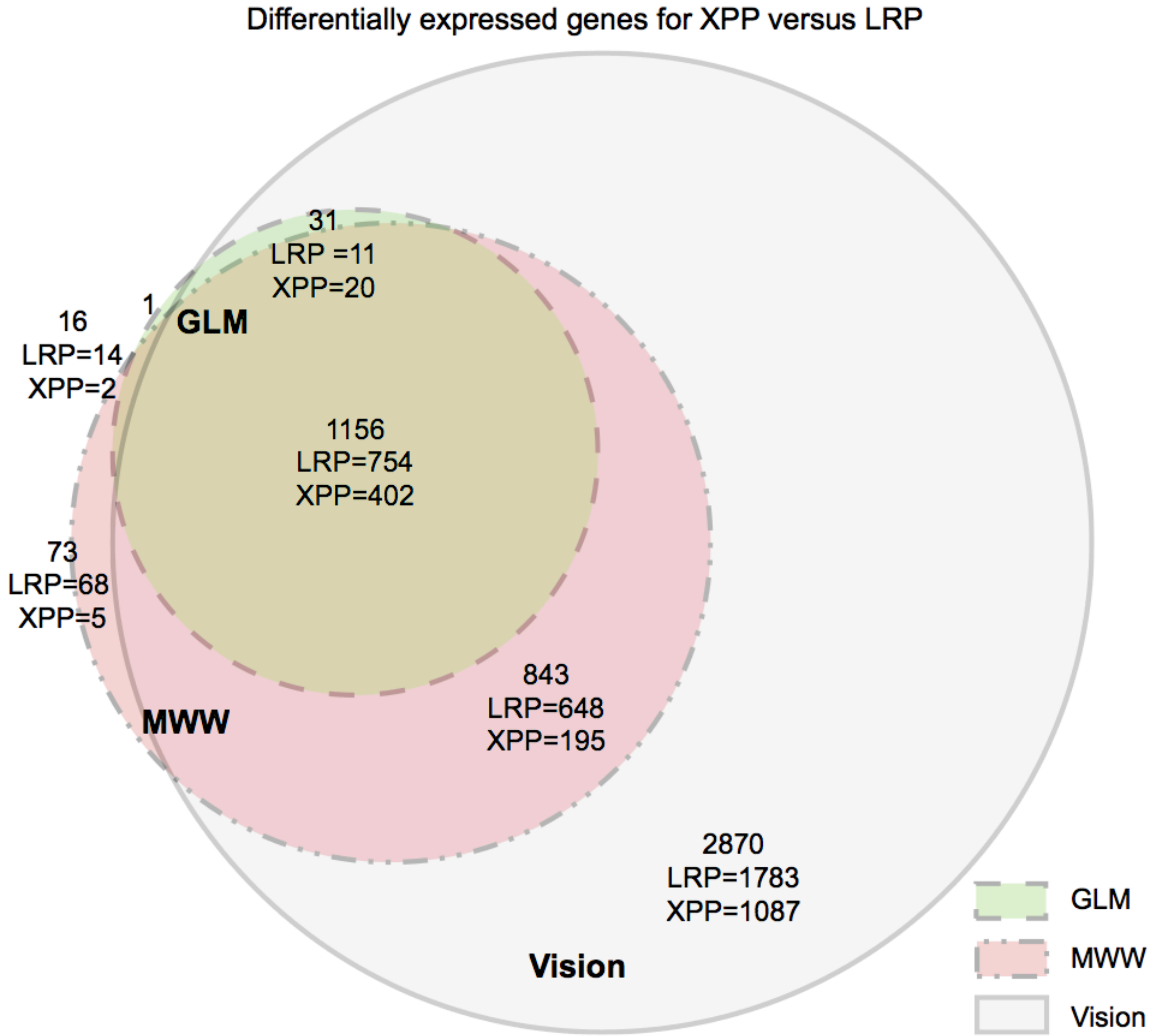


**Supplemental Figure 3. Size-adjusted Venn diagram visualizing overlap of genes between different DEG calling methods.** For each circle of the Venn Diagram, the total number of DEGs as well as the number of DEGs up in LRP and up in XPP are shown. All genes that were called in two or more methods were used for downstream analysis.

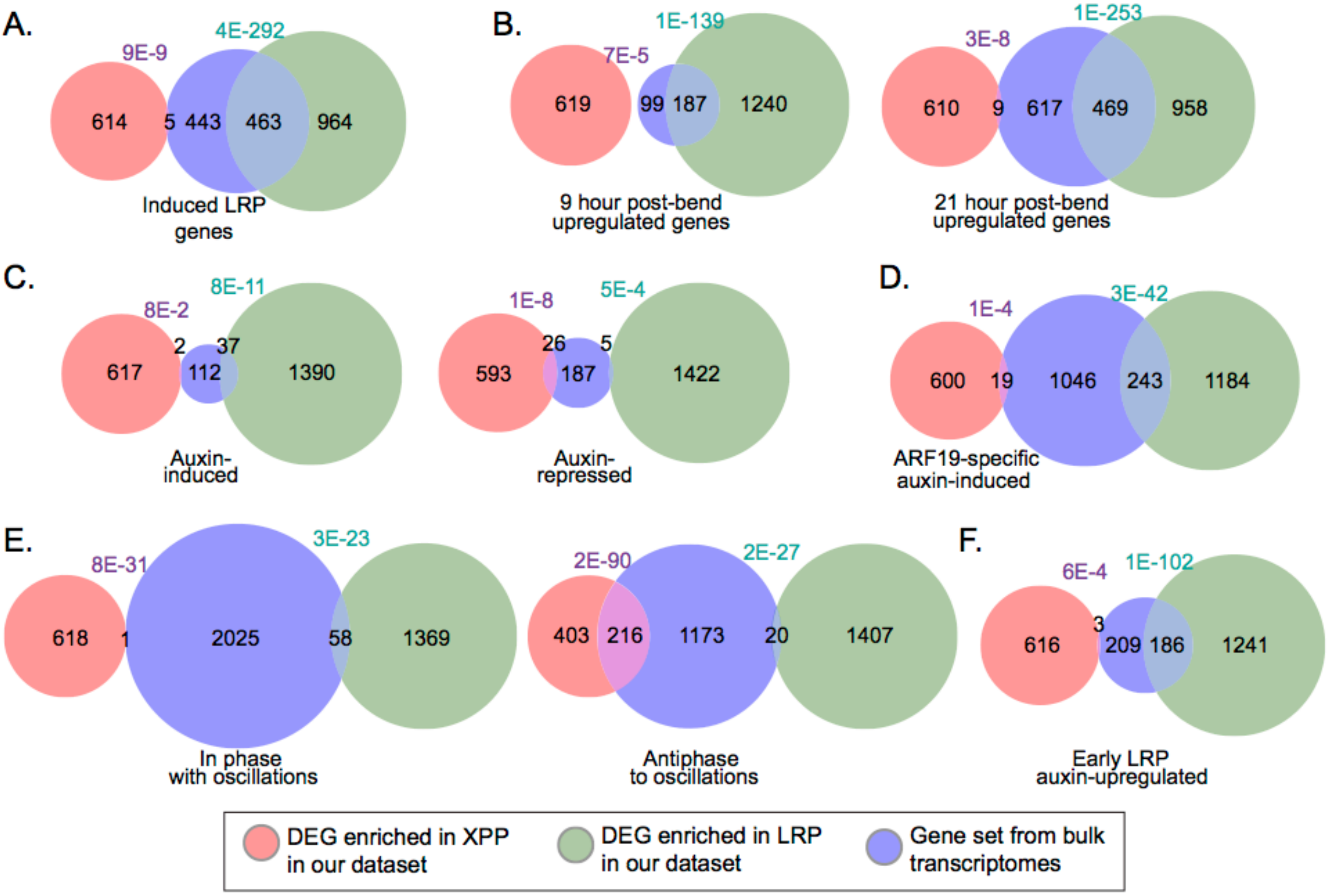


**Supplemental Figure 4. Comparison of XPP and LRP DEGs from the single-cell library to bulk transcriptomes.** A. Comparison to LRP-induced genes from Vanneste et al, 2005. B. Comparison to time-course analysis of lateral root initiation at nine and twenty-one hours post-bend from Voß et al, 2015. C. Comparison to auxin-induced and auxin-repressed genes in the root from Lewis et al, 2013. D. Comparison to ARF19-specific auxin-induced genes from Powers et al, 2019. E. Comparison to genes oscillating in phase and antiphase to auxin in the basal meristem during lateral root specification from Moreno-Risueno et al, 2010. F. Comparison to auxin-induced genes during early lateral root development from Ramakrishna et al, 2019.  Each gene set from bulk transcriptomes is compared to the XPP and LRP DEGs with a size-adjusted Venn diagram. The number of genes in each mutually-exclusive area of each Venn diagram are specified. The values in purple and turquoise denote the hypergeometric distribution for the union XPP/bulk transcriptome and LRP/bulk transcriptome respectively.

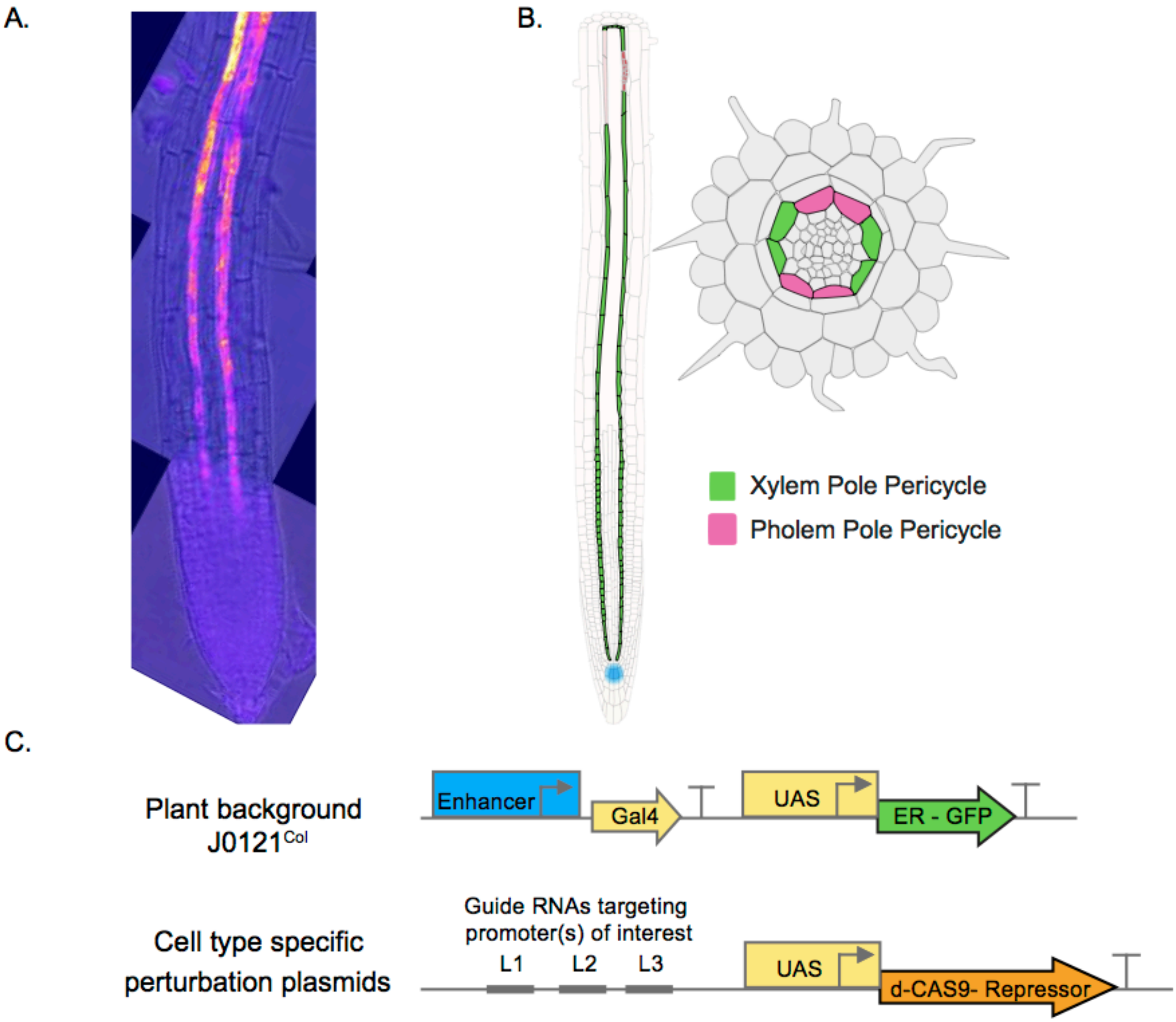


**Supplemental Figure 5. Design of J0121$^{Col}$>>dCas9R system to generate cell type-specific dCas9-repressor mediated knockdown of candidate gene expression.** A. J0121$^{Col}$ is an enhancer trap line where the UAS-Gal4 system drives expression of GFP in the xylem pole pericycle cell file, visualized in confocal microscopy image and B. labeled in cartoons (in green). C. Top panel indicates the enhancer trap cassette in the J0121$^{Col}$ and bottom panels is design of perturbation plasmids included up to three guide RNAs, tagged as location L1, L2, and L3, and a UAS promoter driving expression of dCas9-repressor cassette. Perturbation plasmids with respective cloned guide RNAs were transformed into J0121Col background to drive repression of target genes specifically in xylem pole pericycle cells.

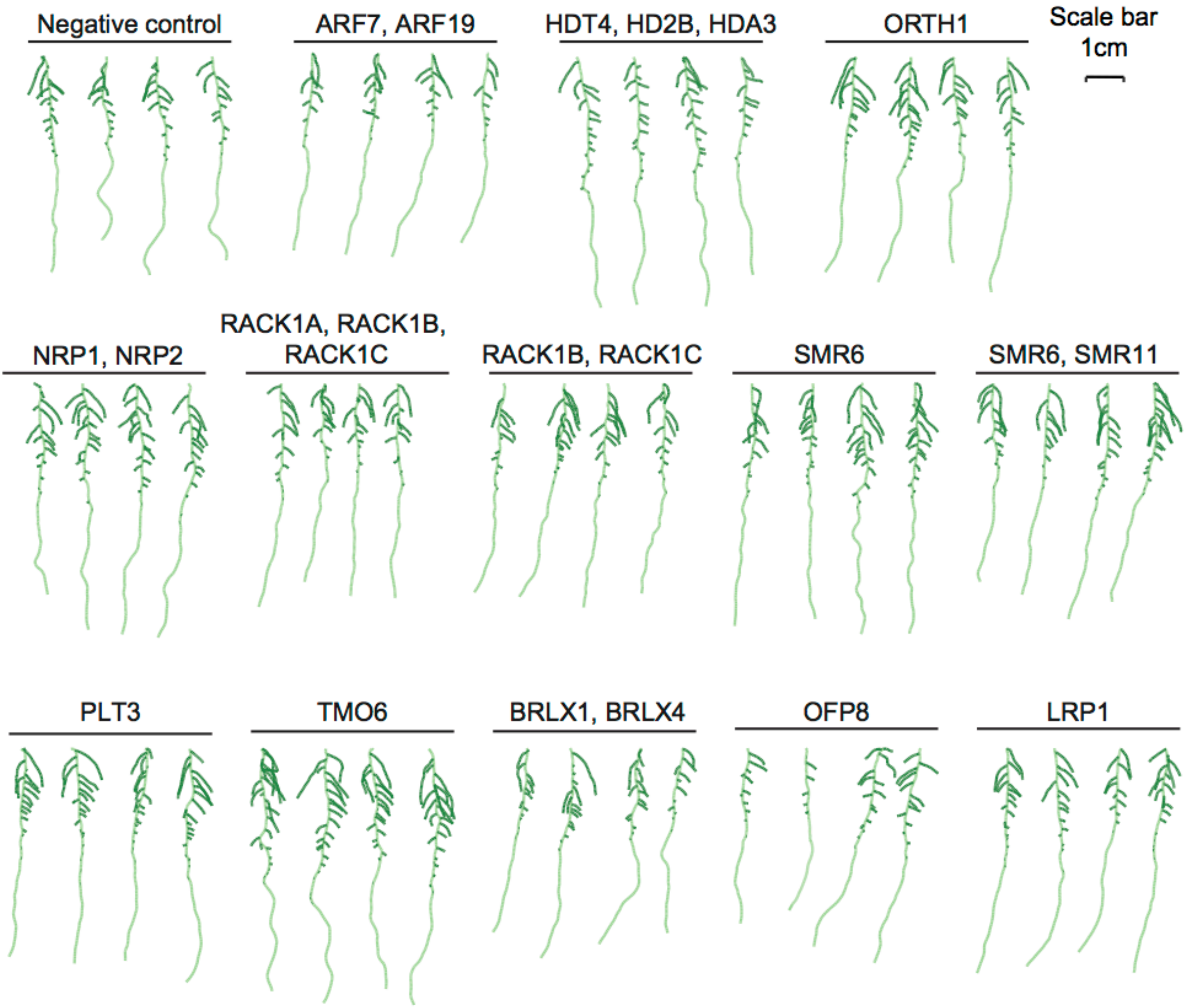


**Supplemental Figure 6. Example seedling traces from J0121$^{Col}$>>dCas9R perturbation lines.** Roots from T2 perturbation line seedlings were quantified for various lateral root developmental phenotypes using SmartRoot.

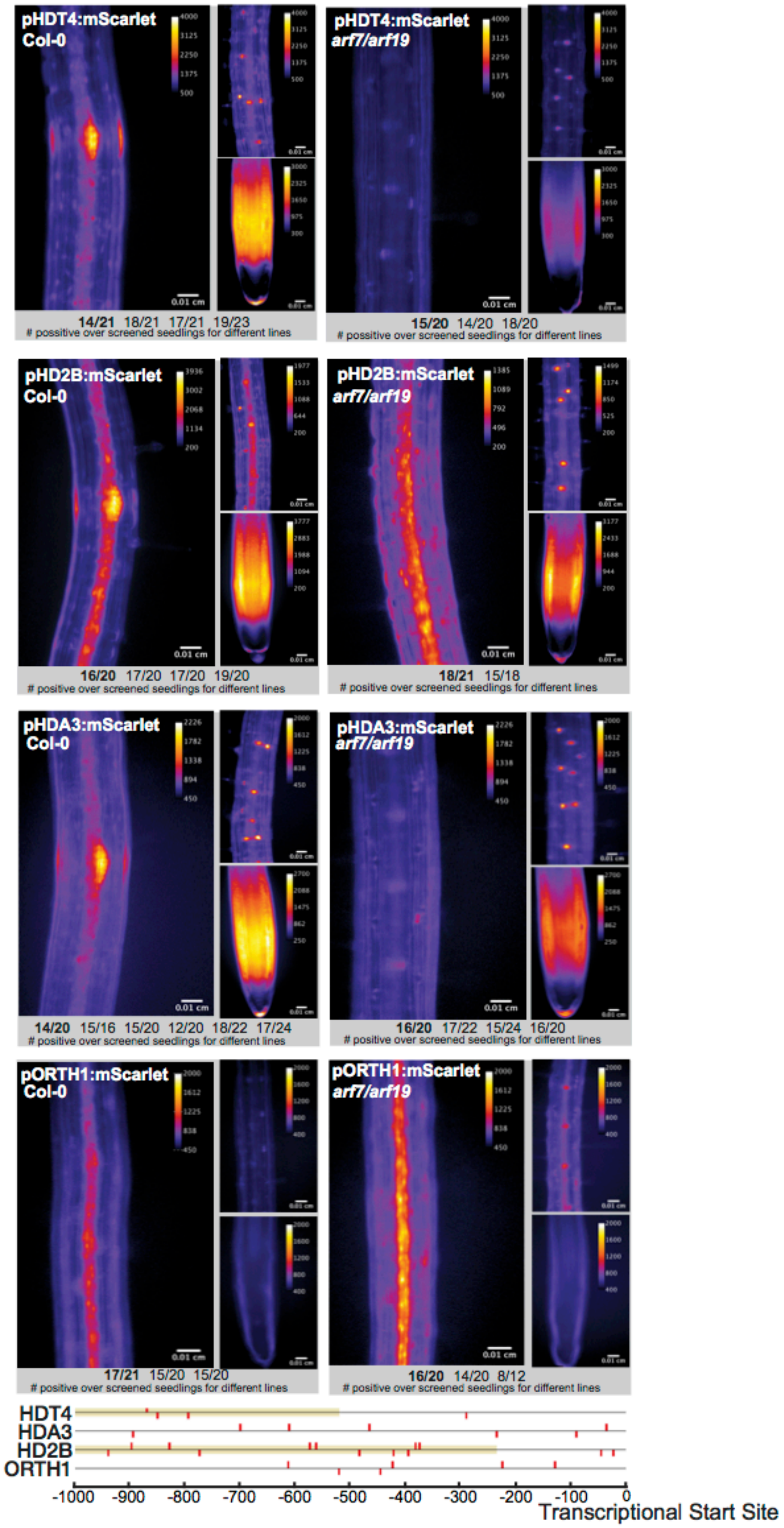


**Supplemental Figure 7. Transcriptional reporters of chromatin regulator candidate genes in wild type and *arf7arf19* roots.** Fluorescent microscopy images of transgenic plant lines carrying transcriptional reporters of candidate genes in early stage lateral root primordia are shown in the large left-side image of each panel. Smaller images of the same transcriptional reporters in a region of the differentiated zone of the root without any developing primordia (above) and in the root apical meristem (below) are shown on the right in each panel. The same reporters in the same regions of the root are imaged in *arf7arf19* mutant background plants on the right. The number of independent transgenic lines imaged per construct and the number of plants within each line that showed expression are reported at the bottom. The lower panel represents 1000 bp upstream of the transcription start site for each gene, with auxin response elements (TGTC/GACA) highlighted in red. Yellow bars indicate CDSs from other genes. This panel was obtained from http://bar.utoronto.ca/cistome.

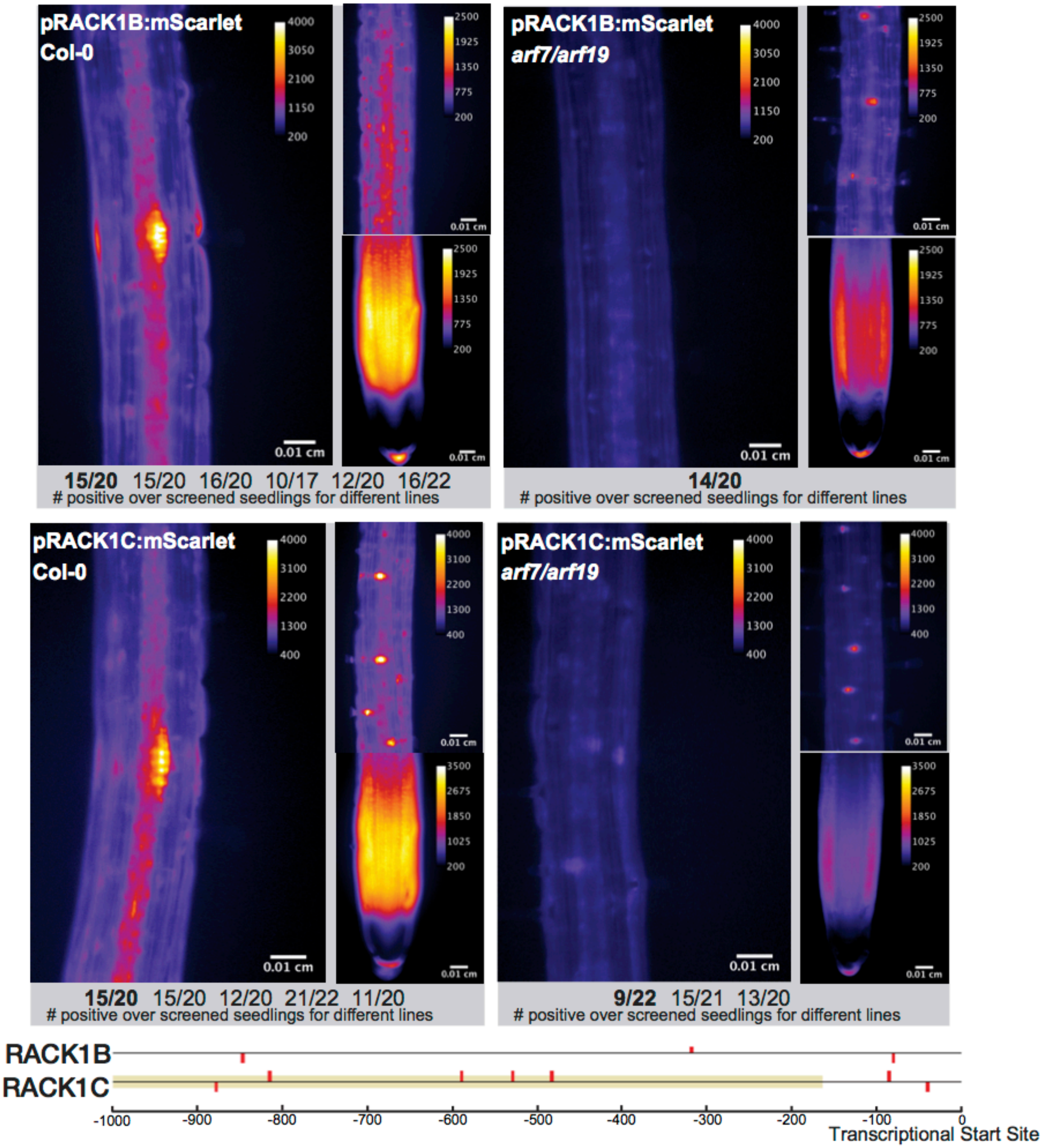


**Supplemental Figure 8. Transcriptional reporters of cell cycle candidate genes in wild type and *arf7arf19* roots.** Fluorescent microscopy images of transgenic plant lines carrying transcriptional reporters of candidate genes in early stage lateral root primordia are shown in the large left-side image of each panel. Smaller images of the same transcriptional reporters in a region of the differentiated zone of the root without any developing primordia (above) and in the root apical meristem (below) are shown on the right in each panel. The same reporters in the same regions of the root are imaged in *arf7arf19* mutant background plants on the right. The number of independent transgenic lines imaged per construct and the number of plants within each line that showed expression are reported at the bottom. The lower panel represents 1000 bp upstream of the transcription start site for each gene, with auxin response elements (TGTC/GACA) highlighted in red. Yellow bars indicate CDSs from other genes. This panel was obtained from http://bar.utoronto.ca/cistome.

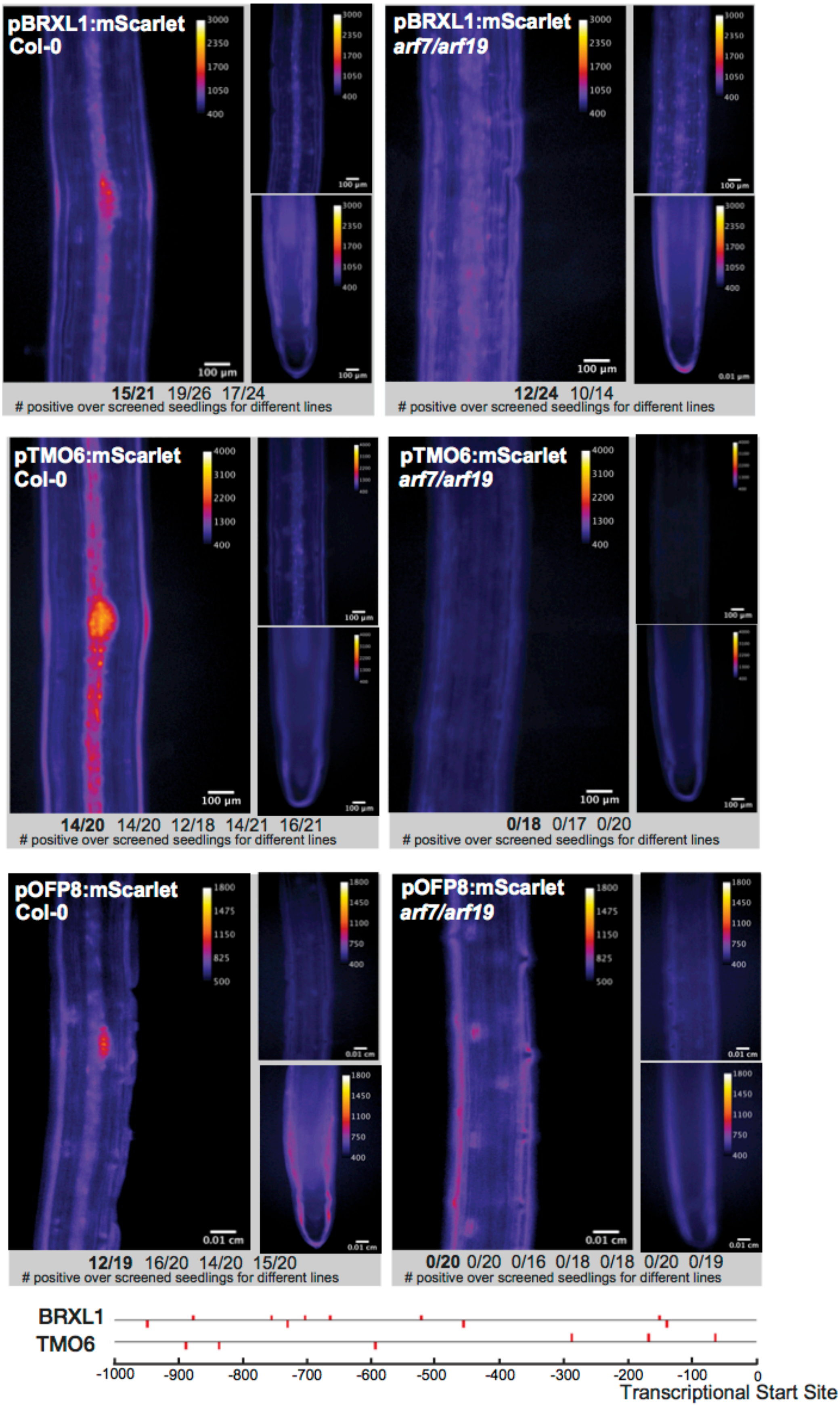


**Supplemental Figure 9. Transcriptional reporters of stemness candidate genes in wild type and *arf7arf19* roots.** Fluorescent microscopy images of transgenic plant lines carrying transcriptional reporters of candidate genes in early stage lateral root primordia are shown in the large left-side image of each panel. Smaller images of the same transcriptional reporters in a region of the differentiated zone of the root without any developing primordia (above) and in the root apical meristem (below) are shown on the right in each panel. The same reporters in the same regions of the root are imaged in *arf7arf19* mutant background plants on the right. The number of independent transgenic lines imaged per construct and the number of plants within each line that showed expression are reported at the bottom. The lower panel represents 1000 bp upstream of the transcription start site for each gene, with auxin response elements (TGTC/GACA) highlighted in red. Yellow bars indicate CDSs from other genes. This panel was obtained from http://bar.utoronto.ca/cistome.

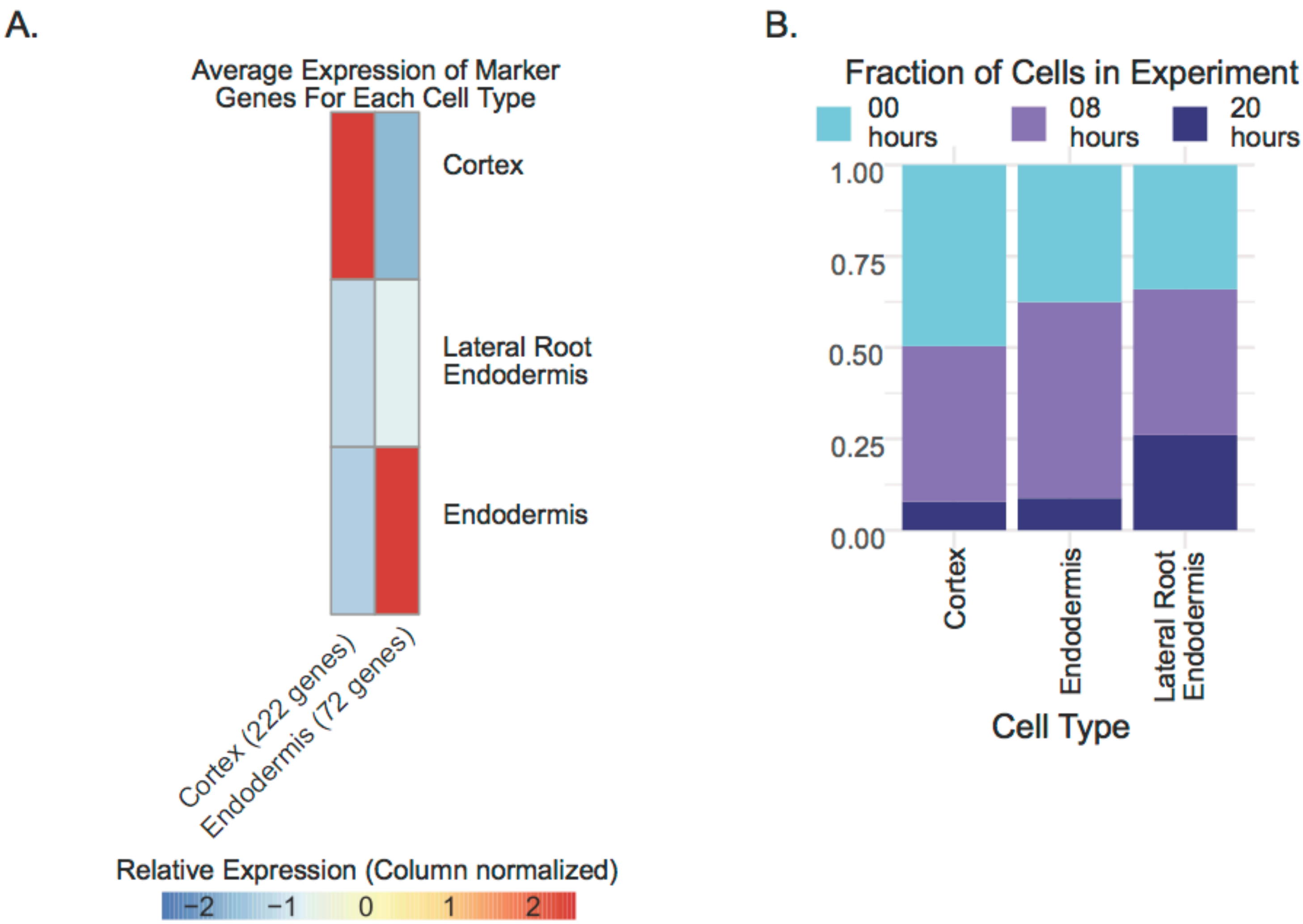


**Supplemental Figure 10. Marker gene expression profiles and experiment breakdown of cortex, endodermis, and lateral root endodermis cells.** A. Heatmap (column-scaled) visualizing average normalized expression of marker genes in the cortex, endodermis, lateral root endodermis (LRE) cells. B. Fraction of cortex, endodermis, and LRE cells from each experiment.

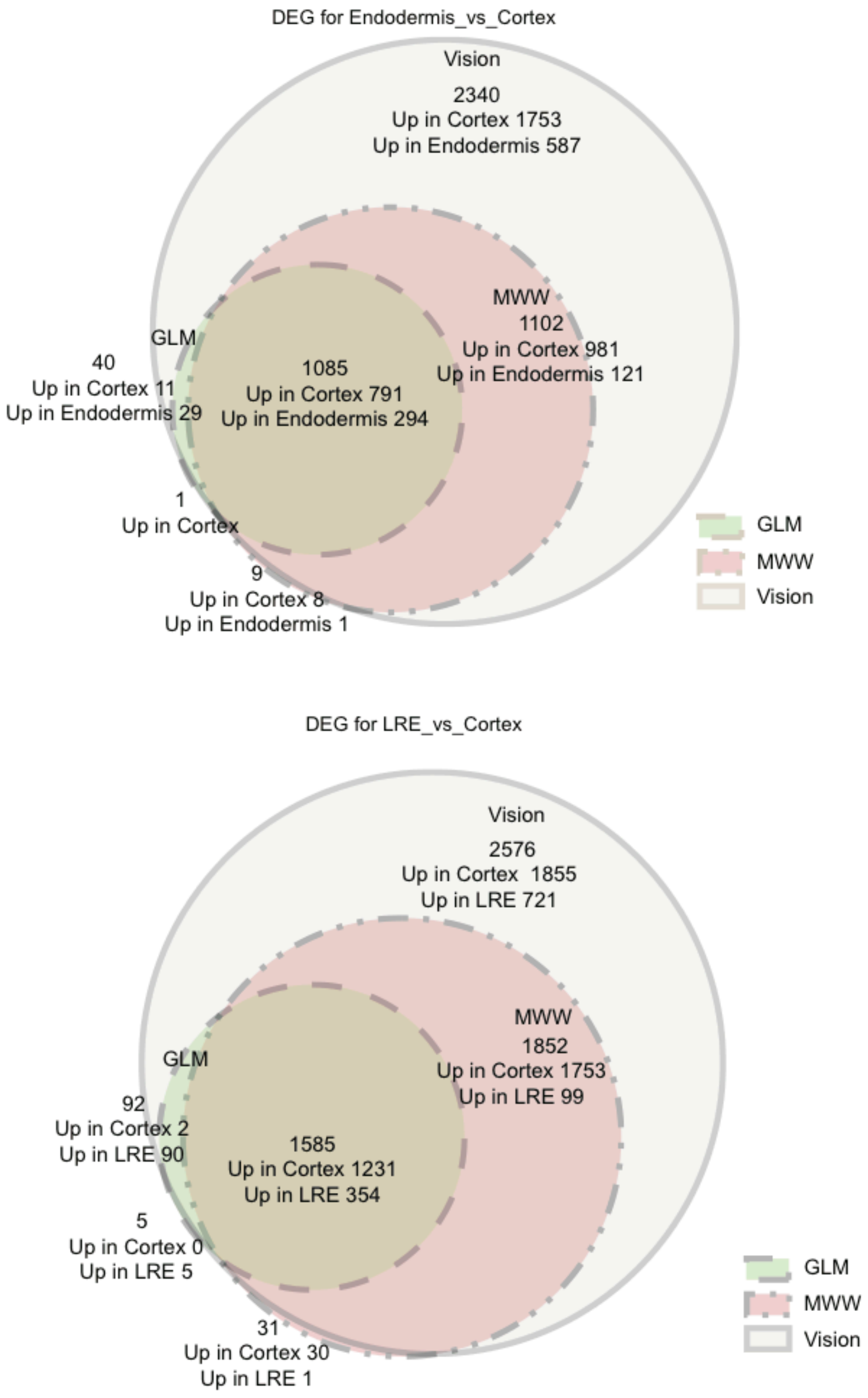


**Supplementary Figure 11. DEG overlaps with different methods for Endodermis/Lateral Root Primodia responding Endodermis analyses** A. Endodermis vs cortex comparison and B. LRE vs cortex comparison. For each ensemble of the Venn Diagram, the total number of DEGs, the number of DEGs up in Cortex and up in Endodermis (for A) and LRE (for B) are added to the diagram.

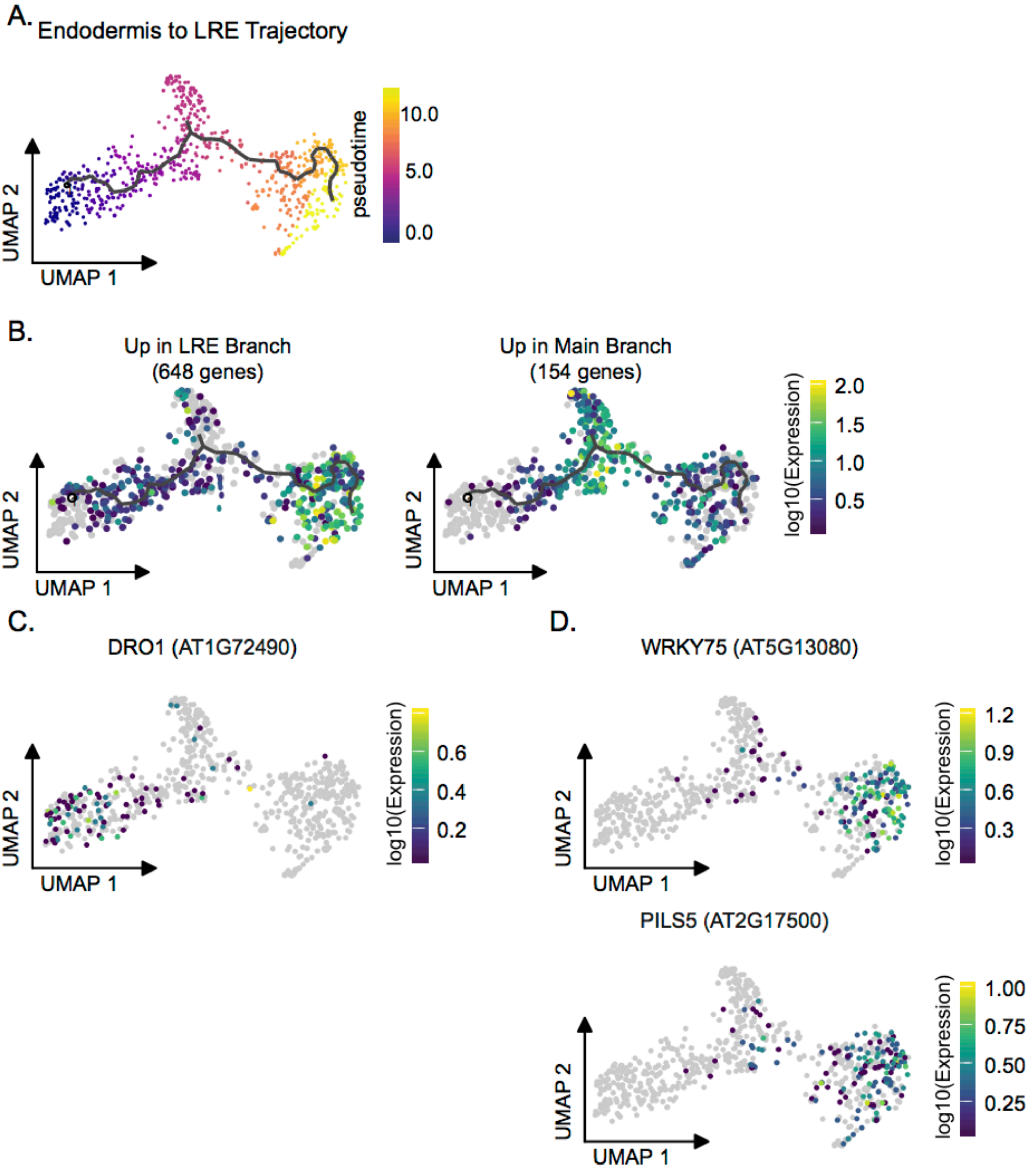


**Supplemental Figure 12. Pseudotime analysis of Endodermis to Lateral Root Endodermis Cells.** A. UMAP of the endodermis to LRE trajectory colored by pseudotime. B. UMAP of the expression of gene sets that differed significantly as a function of pseudotime in the main endodermis branch and the LRE Branch. C. Expression UMAP of DRO1. D. Expression UMAPs of WRKY75 and PILS5

| Cell Type | 00 hour cells | 08 hour cells | 20 hour cells |
|---|---|---|---|
| Columella/ Root Cap | 379 | 207 | 429 |
| Cortex & Endodermis | 311 | 384 | 118 |
| Epidermis | 674 | 545 | 212 |
| Stele | 1121 | 1307 | 971 |
| Total | 2485 | 2443 | 1730 |

| Cell Type | 00 hour cells | 08 hour cells | 20 hour cells |
|---|---|---|---|
| Ambiguous Stele Cells | 212 | 267 | 219 |
| Lateral Root Primordia | 28 | 62 | 77 |
| Mature Pericycle | 92 | 71 | 173 |
| Phloem | 58 | 68 | 90 |
| Phloem Pole Pericycle | 418 | 515 | 273 |
| Xylem | 85 | 102 | 55 |
| Xylem Pole Pericycle | 228 | 222 | 84 |
| Total | 1121 | 1307 | 971 |

| Cell Type | 00 hour cells | 08 hour cells | 20 hour cells |
|---|---|---|---|
| Cortex | 63 | 54 | 10 |
| Endodermis | 154 | 220 | 36 |
| Lateral Root Endodermis | 94 | 110 | 72 |
| Total | 311 | 385 | 108 |

Supplemental Table 1. Breakdown of cell types by experiment
